# Capturing the rhythm: methodological choices shape inference from circadian activity data

**DOI:** 10.64898/2026.10.25.761729

**Authors:** Ashleigh M. Anderson, Kathryn B. McNamara, Theresa M. Jones

## Abstract

1. Statistical tools and analytical decisions for analysing animal activity data are often limited and poorly justified, particularly with respect to model choice, sampling design and temporal aggregation. These shortcomings can constrain our ability to draw accurate and biologically meaningful inferences about behaviour.
2. We interrogated 24 h activity data taken from individual males and females derived from five *Drosophila melanogaster* populations to generate a general framework for collecting and analysing activity data that systematically compares common model classes, continuous verses fragmented sampling strategies, and alternative temporal aggregation choices (time-bin sizes).
3. Our analyses identified generalised additive mixed models (GAMMs) applied to high-frequency continuous data with small-moderate bin sizes (1-5 minutes) as providing the most robust and interpretable inference for our complex, high-frequency activity data.
4. However, our interrogation of the data set confirmed that analytical and sampling decisions need to be tailored to the study species, the timing of activities and the nature of the behaviour of interest. Accordingly, polynomial and cosinor generalised linear models, increased but biologically targeted snapshot sampling, and bin sizes smaller than one hour yielded adequate analyses with similar biological conclusions when appropriately matched to the system.
5. Our approach provides an accessible decision-making framework for activity analyses that allows researchers across behavioural, ecological and evolutionary studies to justify and optimise choices about data collection and modelling. Such an approach improves the biological relevance of inference and supporting the use of modern tracking and analytical technologies.

## Introduction

Circadian activity patterns synchronised to the 24 h day-night cycle are fundamental to life on Earth (Vitaterna et al., 2001). Circadian activity captures the timing of many important biological functions such as foraging, activity, mating and other activities associated with life-histories (Groot, 2014; Vitaterna et al., 2001), with substantial variation among species, sexes and individuals (Tankova et al., 1994). In animals, behavioural changes are often rapid and the first indicators of biotic or abiotic perturbations (Wong & Candolin, 2015), however subtle shifts in activity patterns may be difficult to detect and characterise accurately. Aside from biological differences, a further challenge is that, while activity patterns are widely used to quantify behaviour, analytical approaches vary substantially across, and within, research fields. Methodological decisions made during (i) statistical analysis, (ii) data collection, and (iii) data processing can all influence the representation of activity patterns and potentially their interpretation. These differences can reduce our ability to assess and compare activity patterns across studies and, perhaps more importantly, may mischaracterise the timing, magnitude or shape of activity patterns, potentially leading to inaccurate biological inference.

### Statistical analysis: why are common approaches inadequate?

Activity data are analysed using a wide range of statistical approaches, which vary in their ability to represent complex and non-linear activity patterns. Recent advances in imaging technology, automated tracking, and data storage have provided unprecedented access to high-resolution recordings of animal behaviour across extended time scales (McKenzie-Smith et al., 2025). From continuous video monitoring to sensor-based tracking systems, researchers can now collect vast quantities of behavioural data with fine temporal detail. However, our capacity to statistically process, analyse, and interpret these large and complex datasets has not always kept pace with data acquisition. As a result, much of this information remains underutilised, or is simplified through analytical approaches that fail to capture underlying biological complexity. This mismatch between data availability and analytical capability represents a major bottleneck in contemporary animal behaviour research. Many commonly applied approaches, including generalised linear models (GLM), polynomial regressions and cosinor-based models (Gaston et al., 2013; Koteja et al., 2003; Parsons et al., 2024; Zhang et al., 2011), are largely inadequate at representing circadian activity. These parametric models assume a predefined, rigid and sometimes unrealistic relationship between time and activity, which may not adequately capture complex or irregular activity patterns (Baayen & Linke, 2020; de Freitas et al., 2021; Simpson, 2025). Critically, the oversimplified temporal structure of such models can lead to inaccurate biological interpretation of more complex, temporally longitudinal, activities. Models with more temporal flexibility, such as generalised additive mixed models (GAMMs) and their mixed-effects extensions, address some of these limitations by allowing the shape of the relationship to be determined by the data. This enables the capture of unassumed, non-sinusoidal, multi-peaked dynamics present in activity patterns characteristic of circadian activity (Baayen & Linke, 2020; Parsons et al., 2024; Simpson, 2025; Wood, 2025). Despite their advantages, GAMMS have not been widely adopted across the animal sciences (de Freitas et al., 2021; Simpson, 2025). One reason for this underuse may be that GAMMs are less familiar than traditional modelling approaches, and thus, limited systematic evaluation of their relative performance provides a limited empirical basis for researchers to move away from more established methods.

### Data collection and processing: how do sampling and aggregation decisions alter inference?

Statistical model choice is only one methodological decision that can shape inferences from activity data. Even when an appropriate analytical approach is used, inference can also depend on how activity is sampled and the temporal resolution at which those data are collected and processed. *Sampling* methods for activity data range from high-resolution continuous collection, to low-resolution fragmented approaches, such as snapshot sampling (e.g. 5-min samples every hour for a set period) (Bonilla et al., 2024; Revel et al., 2025). High-resolution continuous sampling methods capture a more complete and detailed record of behavioural dynamics, eliminating the issues of unrepresentative data and distorted activity patterns (Chen et al., 2016; Lynch et al., 2025; Manfrè et al., 2024; McKenzie-Smith et al., 2025; Revel et al., 2025). However, continuous data sampling can generate substantial and often complex datasets (Chen et al., 2016). Consequently, activity data are often simplified either during collection (e.g. through the use of lower resolution, fragmented approaches) or during processing (e.g. by averaging activities across larger time periods).

#### Resolution loss through the sampling window

Fragmented sampling, also known as instantaneous sampling (Manfrè et al., 2024), is common practice in animal behaviour research, likely due to its simplicity and lower computational demands (Chen et al., 2016; Lynch et al., 2025; Manfrè et al., 2024). Fragmented sampling represents a trade-off between sampling speed and temporal resolution. While it can provide an effective means of characterising broad patterns of activity, it may be less efficient at capturing transient behaviours or fine scale temporal variation. Moreover, depending on the sampling regime it may mask or misrepresent activity rhythms or even misclassify inactivity, particularly when detection is spatially constrained (Chen et al., 2016; Lynch et al., 2025; Manfrè et al., 2024), leading to false confidence and errors (Fiske & Delmolino, 2012). Importantly, however, these limitations do not make fragmented sampling inherently inadequate; instead, its ability to accurately represent underlying activity patterns is likely to depend on the specific sampling parameters used. Where fragmented sampling is used, two parameters are likely of most importance: the duration and timing of the sampling window (Brereton et al., 2022; Chen et al., 2016; Lynch et al., 2025; Manfrè et al., 2024). A longer sampling window increases the likelihood of capturing transient behaviours, in addition to more closely approximating patterns derived from more continuous data sets. Sampling timing is equally, if not more, important, as activity is typically not uniformly distributed across any given temporal period (e.g. hour, day, season). As a consequence, the timing of the snapshot schedule (regularly spaced random sampling vs. targeted biologically relevant sampling) may capture different components of an activity rhythm yielding different biological conclusions. Thus, both the duration and timing of the sampling window can influence how accurately fragmented data represent biological reality which may have consequences for broader interpretation.

#### Resolution loss during data processing

Regardless of the sampling approach, activity monitoring can generate substantial, complex datasets that require some degree of simplification prior to formal analysis (Lynch et al., 2025; Simpson, 2025). Simplification often occurs through the aggregation of data into discrete time bins which accelerates computation time, reduces noise and can facilitate biologically meaningful statistical analyses (Dowse, 2009; Rajaguru et al., 2025). Resulting time bins can range from a few seconds up to 24 h depending on the temporal scale and duration of the original sampling period. However, as with fragmented sampling, the choice of bin size represents a trade-off between retaining adequate temporal information and reducing data complexity (Dowse, 2009; Dowse & Ringo, 1994). Smaller bin sizes preserve finer scale variation, but they may also produce noisier, less interpretable, activity profiles Dowse, 1994 #587;Rajaguru, 2025 #676}. In contrast, larger bin sizes may obscure short term shifts in behaviours or smooth otherwise distinct activity peaks (Dowse & Ringo, 1994; Langener et al., 2024; Rajaguru et al., 2025). The appropriate level of aggregation (e.g. bin size) again depends on the temporal scale of the activity of interest (e.g. short periodic bursts v longer duration behaviours) and the resolution of the original data set (Dowse, 2009; Dowse & Ringo, 1994). Despite the importance of these decisions, there is little standardisation in sampling regimes or binning strategies, within and across species and research fields, potentially limiting the accuracy of activity estimates and biological interpretations, as well as cross-study comparisons.

The above overview highlights several important knowledge gaps with respect to analysis and interpretation of activity data. First, the relative ability of commonly used statistical approaches to represent complex activity patterns has not been widely systematically evaluated, making it unclear when simpler parametric models are adequate and when more flexible approaches are required. Second, although fragmented sampling is widely used and may be the only feasible logistic option, we have limited understanding of how the duration and timing of sampling windows influence the representation of underlying activity patterns, or the extent to which fragmented sampling can approximate continuous data collection. Similarly, there is little empirical guidance regarding the level of temporal aggregation that can be applied to high-resolution data before biologically meaningful information is lost.

To address these challenges, we adopt a case study approach using the vinegar fly, *Drosophila melanogaster*, a well-established model organism in circadian and behavioural research (Allada & Chung, 2010)*. Drosophila melanogaster* has robust and well-characterised daily rhythms, distinct locomotor activity patterns, and well-defined periods of rest (Chiu et al., 2010; Flatt, 2020) providing a model biological system for evaluating how methodological decisions influence biological interpretation of activity. Critically, there are a number of existing data sets and analytical approaches for *Drosophila* providing relevant comparators for our approaches (Chiu et al., 2010; Revel et al., 2025; Zhang et al., 2011).

To systematically evaluate the three methodological decisions outlined above - statistical analysis, data collection and data processing - we used a large circadian activity data set of 114 24 h high-resolution continuous video recordings across five genetically distinct *D. melanogaster* populations. Specifically, we compared (1) the ability of commonly used statistical model-types and GAMMs to represent underlying activity patterns, (2) the effects of sampling window duration and timing by comparing snapshot data with continuous data collection; and, (3) the effects of temporal aggregation (bin size) on the representation and biological interpretations of activity patterns. Our overall aim was to assess how these decisions made during collection, processing and analysis affect biological inference and to use these findings to generate an evidence-based framework for the accurate and standardised analysis of animal activity data.

## Materials and Methods

### 3.1 Study system and activity recordings

Experimental *D. melanogaster* populations were obtained from the Bloomington *Drosophila* Stock Centre (Indiana, USA). To evaluate the methodological approaches across a range of potential underlying activity patterns, we selected five inbred populations from the *Drosophila Genetic Reference Panel* (DGRP) known to differ in their circadian activity phenotypes (Harbison et al., 2019) (Table S1). Flies were reared under standardised common-garden conditions for four generations prior to experimentation (12 h light:12 h dark cycle; 25°C; ∼65% humidity; *ad libitum* diet of Bloomington’s cornmeal food recipe supplemented with live yeast; see Supplementary Materials). Individuals were video monitored through eclosion to establish precise adult age. Activity was subsequently quantified over a 24 h period beginning at ZT 0, with individuals aged 4-28 h at the start of recording (see Supplementary Materials).

Activity was recorded using continuous high-resolution video monitoring under controlled photoperiod conditions (12 h light:12 h dark). Video recordings captured both photophase (day) and scotophase (night) behaviours and were processed using EthoVision XT (Version 18; Noldus Information Technology, Wageningen, The Netherlands). Further details regarding animal husbandry, recording setup, and video processing are provided in the Supplementary Materials.

### 3.2 Behavioural metrics

From the continuous recordings, we extracted four complementary measures of activity aggregated in 1-min bins using EthoVision XT: (i) total distance moved, (ii) time spent moving, (iii) movement bout frequency and (iv) rest behaviour. Together, these variables capture complementary dimensions of locomotor activity and circadian organisation commonly quantified in behavioural and chronobiological studies (Table 1).

**Table 1.**
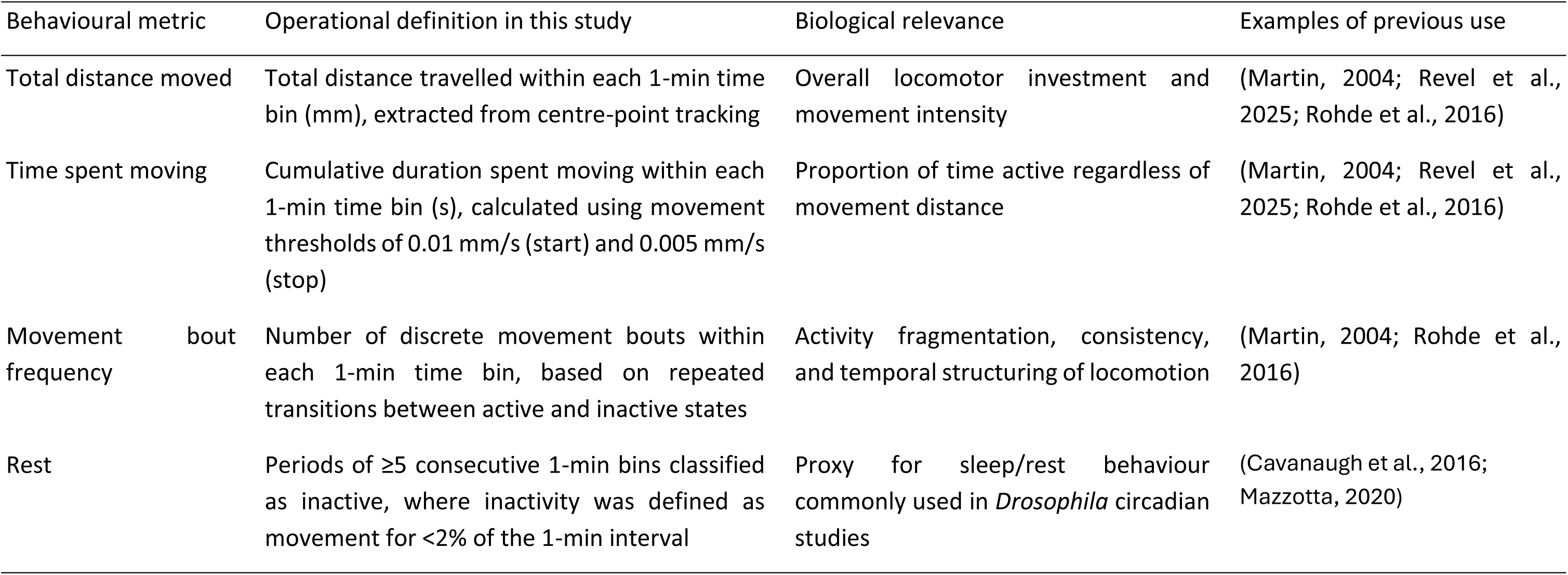
Behavioural metrics extracted from continuous activity recordings, their operational definitions in this study, biological relevance and examples of their previous use in behavioural research.

Rest behaviour was used as a proxy for sleep and was defined as ≥5 consecutive min of inactivity from the aggregated data in 1-min bins, consistent with standard *Drosophila* sleep literature (Cavanaugh et al., 2016; Mazzotta, 2020). Full behavioural extraction settings and threshold definitions are provided in the Supplementary Materials.

### 3.3 Evaluating methodological approaches for activity data

We evaluated one key methodological decision at each of the three stages identified above: (i) statistical model choice, (ii) sampling strategy, (iii) temporal aggregation (bin size).

#### 3.3.1 Comparing the efficacy of statistical modelling approaches (using 1-min bins)

We compared four commonly used model classes: (i) linear models, (ii) quartic polynomial models, (iii) GLMM-cosinor models, and (iv) GAMMs. For each behavioural metric (total distance moved, time spent moving, movement bout frequency, and rest), we fitted: a linear model, two variants of quartic polynomial models, two variants of a GLMM-cosinor model, and a first order autoregressive residual (AR1) GAMM.

Separate variants of the polynomial and GLMM-cosinor models were used for statistical comparison and visualisation. For the polynomial analyses, we fitted both orthogonal quartic polynomial models and raw quartic polynomial models. Orthogonal polynomials were used for statistical comparisons because they reduce multicollinearity among polynomial terms. Raw polynomials were used for visualisation because they retain the original temporal scale (24 h represented by 1,440 1-min intervals) facilitating visual interpretation of activity patterns across the daily cycles.

For the GLMM-cosinor analyses, we fitted models over a 12 h or 24 h periodicity. The 24 h models capture the underlying circadian periodicity allowing estimation of circadian amplitude and acrophase. The 12 h models were fitted to provide a closer visual representation of the characteristic bimodal morning and evening activity peaks seen in *D. melanogaster* (Pegoraro et al., 2020; Shaw et al., 2019). Accordingly, the 12 h models were adopted primarily for visual inspection across modelling approaches rather than explicit statistical inference.

#### 3.3.2 Comparing the efficacy of continuous v snapshot sampling

We used a two-step approach to assess how the timing and duration of fragmented (snapshot) sampling influenced the representation of circadian activity patterns. For both comparisons, snapshot datasets were generated from the continuous 24-h recordings, with activity summed within each designated observation window to produce a single value per individual per snapshot.

First, we evaluated the effect of sampling timing while holding sampling duration constant at 60 min. We generated two piecewise continuous snapshot sampling schemes (Lynch et al., 2025), each comprising six 60-min observation windows distributed across the 24-h cycle, with activity summed within each window to produce a single observation per individual. The biologically informed scheme used observation windows associated with major circadian transitions and expected activity peaks or troughs in *D. melanogaster* activity (Pegoraro et al., 2020; Shaw et al., 2019). We used 60-min observation windows beginning at Zeitgeber Time (ZT) 0 (lights-on), ZT 6 (midday), ZT 11 (1 h before lights-off), ZT 12 (lights-off), ZT 18 (midnight), and ZT 23 (1 h before lights-on). The second scheme used six evenly spaced 60-min observation windows beginning at ZT 2, ZT 6, ZT 10, ZT 14, ZT 18 and ZT 22. Comparison of these two schemes allowed us to determine whether biologically informed sampling more accurately represented continuous activity patterns than evenly spaced sampling.

Second, we evaluated the effect of sampling duration while holding the biologically informed sampling schedule constant. At each of the six biologically informed sampling times, we extracted a single 5-, 15-, or 60-min observation window beginning at the nominated ZT, and summed activity within that window to produce one observation per individual. We then compared the activity patterns derived from each snapshot duration with those derived from the continuous reference dataset. This allowed us to determine how reducing the duration of observation at biologically relevant times influenced the ability of fragmented sampling to represent the underlying continuous activity pattern.

#### 3.3.3 Temporal aggregation and bin-size comparison

To assess how increasing temporal aggregation influenced inferences about activity patterns, we progressively aggregated the high-resolution, continuous data set into coarser temporal bins of 1, 5, 15, 30, 60 min, and 12 h. Starting from an existing fine-scale time index (1-min bins), we defined new, non-overlapping time bins of the desired width by grouping the neighbouring minutes. Within each bin, and for each individual, we summed the four behavioural measures, yielding one aggregated record per individual per time bin. This approach allowed us to assess the effects of temporal aggregation (bin size) while holding the underlying activity data constant.

Data from the 1- to 60-min bins were analysed using a GAMM structure. Aggregation into 12 h bins reduced the temporal structure to only two observations per circadian cycle, providing insufficient temporal information to reliably fit a GAMM. We therefore used generalised linear mixed models for the 12-hr bins. Details of the model specifications are provided in Section 3.4.3.

### 3.4 Data Analysis

All analyses were conducted using R (version 4.6.1). Individuals missing >25% of tracking points (likely resulting from video contrast or tracking errors) were excluded from analyses (N = 25). Missing bins were treated as missing rows with no imputation. The final dataset comprised of video files from 114 flies (between 22-24 flies per replicate population) (Table S2).

As *D. melanogaster* exhibit sexually dimorphic activity patterns (Asahina, 2018), all analyses were conducted separately for males and females. All models included terms for time, wing size (Supplementary materials; Section 1.4), and age (hours since eclosion at start of recording; Supplementary materials, Section 1.3) to account for temporal changes in activity, size-related variation in movement, and age-related behavioural shifts (Giebultowicz & Long, 2015; McKenzie-Smith et al., 2025), respectively. Wing size and age were included as covariates to account for potential sources of variation but were not themselves the focus of biological inference. Replicate population and individual identity were included as random effects to account for population-level variation and repeated observations across time.

All GAMMs, linear and polynomial models used in the model-class comparisons were fitted using bam() in the mgcv package (Wood, 2017) with fast Restricted Maximum Likelihood (fREML) estimation to improve computational efficiency given the size of our time-series datasets (Wood, 2017). GLMM-cosinor models were fitted using the GLMMcosinor package (Parsons et al., 2024). See Supplementary Materials for a more detailed overview of the modelling setups (Section 2.1).

For GAMMs, temporal smooths were fitted using penalised cubic regression splines with shrinkage (bs = "cs") to limit overfitting. GAMM smooths were visualised using the smooth_estimates() function from the gratia package (Simpson, 2024) and predicted activity curves were plotted using ggplot2. We chose an AR1 GAMM structure using the itsadug package. The AR1 structure was used to account for autocorrelation in model residuals (Wood, 2017). The maximum basis dimension (k-value) was determined for each model to ensure models maintained sufficient flexibility while reducing overfitting (model-specific values are provided in Table S3). Basis adequacy was assessed using gam.check() and AIC comparisons across models with varying maximum basis dimensions. The prediction uncertainty (95% confidence intervals) was calculated on the link scale and back transformed for linear, polynomial, and GAMM models. Confidence intervals (95%) were used for the GLMM-cosinor models. Residual diagnostics and overdispersion checks for GAMMs and linear models were conducted using the DHARMa package.

All GAMMs, linear and polynomial models were fit with tweedie log-link distributions for total distance moved and time spent moving, negative binomial log-link models were used for movement bout frequency, and binomial logit-link models were used for rest behaviour to account for zero inflation and overdispersion of count structures in the behavioural data (de Freitas et al., 2021; Simpson, 2025; Wood, 2017). For the GLMM-cosinor models, we used a gamma log-link for total distance moved and time spent moving (with the addition of a small positive constant (+ 1e-6) to avoid zero values for model fitting).

#### 3.4.1 Comparing the efficacy of various modelling approaches

To compare efficacy of the various model classes, we assessed model performance using Akaike Information Criterion (AIC), deviance explained, R² values, and visual comparison of predicted activity curves relative to the observed data for the GAMMs, linear and polynomial models. (Bera et al., 2021; Simpson, 2025; Wood, 2017). Combined, these measures allowed us to compare the performance of our statistical models and the ability of each model to represent the observed temporal structure of a specific activity. For the GLMM-cosinors, we converted the estimated acrophase metric (the timing of the maximum predicted activity) into ZT to allow comparison of peak activity timing with that predicted by the GAMMs.

#### 3.4.2 Comparing the efficacy of continuous v snapshot sampling

We used an AR1 GAMM structure for the continuous and various snapshot datasets. As these models differed substantially in both dataset size and temporal structure, direct comparisons using AIC, deviance explained, or fREML scores were not appropriate. Instead, we used each snapshot model to predict activity across the full 24-h cycle and compared the resulting activity curves with predictions from the full 5-min continuous dataset. Predictive performance was assessed using unscaled Root Mean Square Error (RMSE), R² scores (Baayen & Linke, 2020; Bera et al., 2021; Simpson, 2025) and visual comparison of the activity patterns.

#### 3.4.3 Temporal aggregation and bin-size comparison

Here, our aim was to identify which aggregation interval (bin size) best preserved the underlying temporal pattern, rather than to compare absolute goodness-of-fit across differently scaled responses. We therefore treated the highest-resolution model (1-min bins) as our reference point and, for each behaviour and sex, compared the corresponding lower-resolution models (5-, 15-, 30- and 60-min bins) by aligning their predicted curves in Zeitgeber time. We then calculated RMSE and R² on z-standardised predictions (Wermelinger et al., 2025), together with the correlation between curves. These metrics quantified how closely each of the lower resolution models reproduced the shape of the fine-scale 1-min reference activity pattern, independent of differences arising from aggregation. The 12 h bin models were assessed separately by examining model estimates and test statistics for the time, wing and age variables, together with visual comparison of the activity patterns.

## Results

As the four behavioural metrics led to qualitatively similar conclusions across the methodological comparisons, we present time spent moving as a representative behavioural metric in the main text. Results for total distance moved, movement bout frequency, and rest behaviour are provided in the Supplementary Materials (Section 3). Age and wing size were included as covariates in all models but were not the focus of the model comparison; their parameter estimates and statistical tests are reported in the Supplementary Materials (Section 3).

### 4.1 Comparing the efficacy of various modelling approaches

For the time spent moving, GAMMS provided the closest visual representation of the observed temporal activity pattern in both females and males (Table 3). Statistically, GAMMs also showed the strongest model performance, compared to the linear and polynomial approaches, with lower AIC values and higher deviance explained and *R^2^* values (Table 3a).

**Table 3A.** Model-performance metrics for linear, quartic polynomial and generalised additive mixed models (GAMMs) fitted to time spent moving in females and males. Bold values indicate the strongest performance within each sex and metric. GLMM-cosinor models are presented separately because these model-performance metrics were not used for direct comparison with the other model classes.

| Model | Female |  |  | Male |  |  |
| --- | --- | --- | --- | --- | --- | --- |
|  | AIC | R<sup>2</sup> | Deviance explained (%) | AIC | R<sup>2</sup> | Deviance explained (%) |
| Linear | 349862.1 | 0.19 | 17.0 | 226221.8 | 0.09 | 11.7 |
| Quartic polynomial | 346987.8 | 0.23 | 20.5 | 223378.3 | 0.12 | 17.1 |
| GAMM | **304178.5** | **0.25** | **23.2** | **178571.1** | **0.16** | **24.1** |

All four model classes detected significant temporal variation in activity but differed in their capacity to represent its temporal structure (Table 3). The polynomial and GLMM-cosinor models captured the broad shape and temporal pattern of the activity, but tended to over- or underestimate the magnitude of the activity peaks relative to the observed data (Fig. 1). In contrast, the linear model, provided a poor representation of the nonlinear activity pattern (Fig. 1).

**Figure 1.**
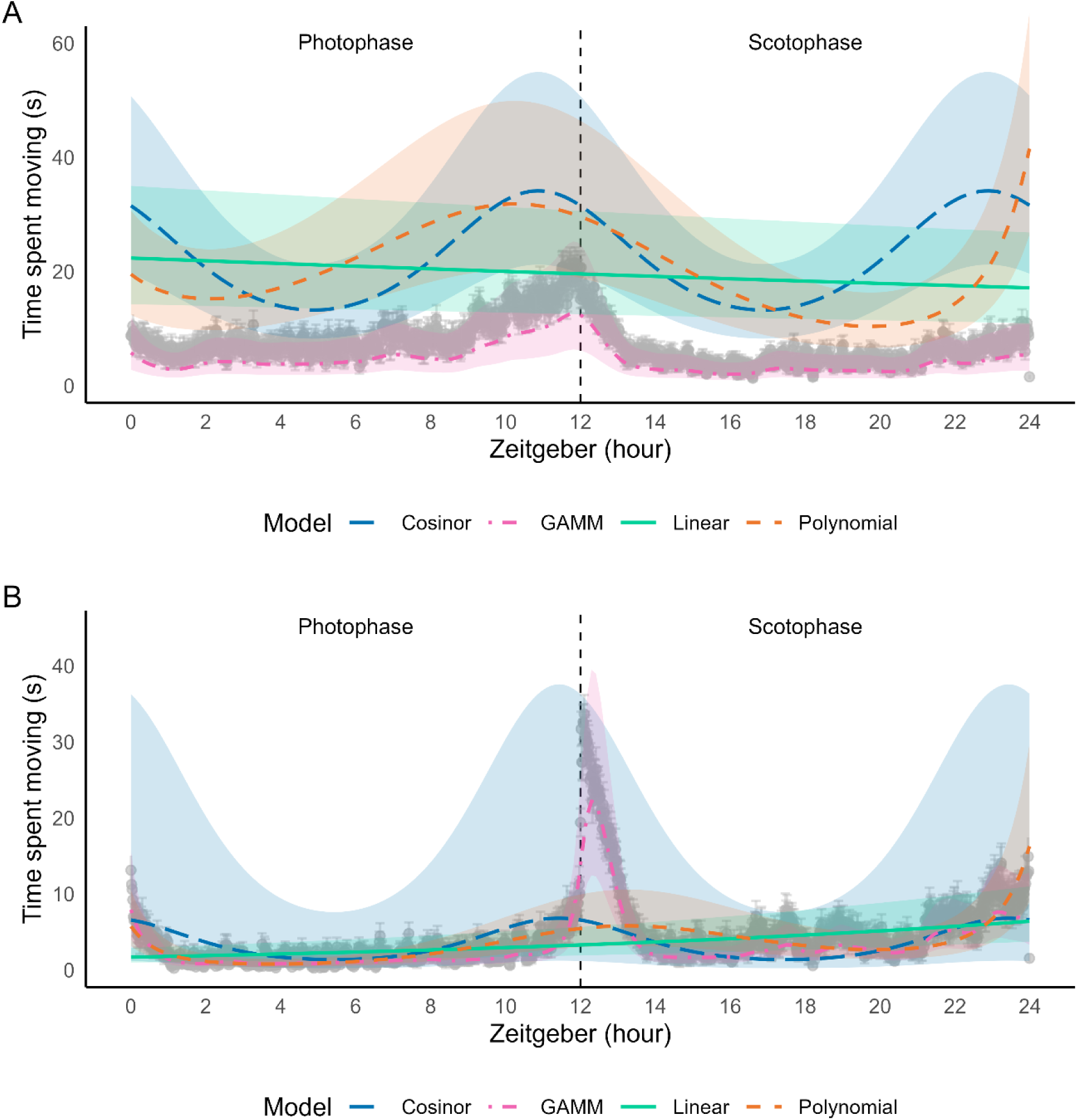
Comparison of observed and model-predicted time spent moving over 24 h for (A) females and (B) males, using data aggregated into 1-min bins. Raw observed data is shown in grey (± standard errors); lines show predictions from the linear n solid), quartic polynomial (orange dashed), GLM-cosinor (blue dashed), and GAMM models (pink dashed).

Estimates of the timing of peak activity also differed across the modelling approaches. GAMMs identified the activity peak around ZT 12 (lights-off) in both males and females (Fig. 1). For females, the GLMM-cosinor model encompassed this peak, with estimated peak activity between ZT 11.72 – 14.34. For males, the corresponding estimate was substantially less precise, ranging between ZT 7.07 and 20.33 (Table 3b).

**Table 3B.**
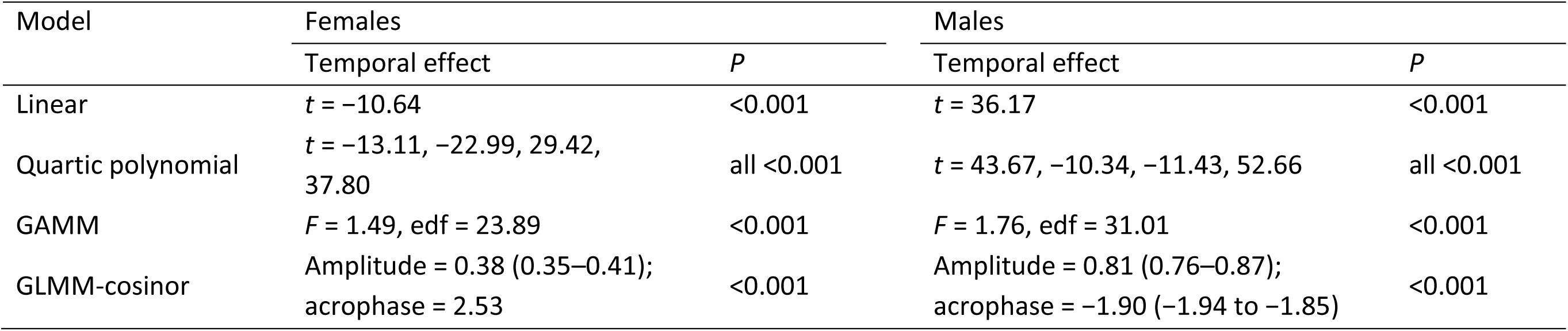
Temporal effects on time spent moving in females and males estimated using linear, quartic polynomial, generalised additive mixed models (GAMMs) and GLMM-cosinor models. For polynomial models, *t*-statistics are shown for each of the four polynomial terms. GAMM results report the *F*-statistic and estimated degrees of freedom (edf). GLMM-cosinor results report amplitude and acrophase estimates with 95% confidence intervals in parentheses. Age and wing size were included as covariates in all models but were not the focus of the model comparison; their parameter estimates and statistical tests are reported in Table S4.

| Model | Females |  | Males |  |
| --- | --- | --- | --- | --- |
|  | Temporal effect | *P* | Temporal effect | *P* |
| Linear | *t* = -10.64 | <0.001 | *t* = 36.17 | <0.001 |
| Quartic polynomial | *t* = -13.11, -22.99, 29.42, 37.80 | all <0.001 | *t* = 43.67, -10.34, -11.43, 52.66 | all <0.001 |
| GAMM | *F* = 1.49, edf = 23.89 | <0.001 | *F* = 1.76, edf = 31.01 | <0.001 |
| GLMM-cosinor | Amplitude = 0.38 (0.35–0.41);<br>acrophase = 2.53 | <0.001 | Amplitude = 0.81 (0.76–0.87);<br>acrophase = -1.90 (-1.94 to -1.85) | <0.001 |

### 4.2 Comparing the efficacy of continuous v snapshot sampling

Focusing on time spent moving as the representative behavioural metric, we first assessed the effect of snapshot timing by comparing 60-min biologically informed and evenly spaced sampling windows. For both females and males, biologically informed snapshots (ZT0, 6, 11, 12, 18 and 23) more closely reproduced the continuous activity patterns than evenly spaced snapshots (ZT2, 6, 10, 14, 18 and 22). Biologically informed snapshots had lower RMSE and higher R² values relative to the continuous reference dataset, indicating greater similarity to activity patterns derived from continuous sampling (Table 4).

**Table 4.** The Root Mean Square Error (RMSE) and *R*^2^ scores for the predicted continuous data from each of the four snapshot models (5-min, 15-min, 60-min, and 60-min of evenly spaced snapshots) compared to the continuous 5-min data for the time spent moving behaviour. The values that indicate the best fit compared to the continuous data are in bold.

| Model | Females |  | Males |  |
| --- | --- | --- | --- | --- |
| | RMSE | $R^2$ | RMSE | $R^2$ |
| 5-min | 0.395 | 0.162 | 0.745 | 0.121 |
| 15-min | 0.274 | 0.597 | 0.715 | 0.189 |
| 60-min | **0.256** | **0.647** | **0.519** | **0.572** |
| 60-min evenly spaced | 0.273 | 0.601 | 0.638 | 0.354 |

Visual comparison of the GAMM temporal smooths revealed some sex-specific differences in the effect of snapshot timing. For females, both 60-min sampling schemes reproduced the broad shape of the continuous activity pattern; however, evenly spaced snapshot sampling failed to accurately reproduce the major activity peak at ZT12 (Fig. 2). For males, the effect was more pronounced, with evenly spaced snapshots providing a poorer representation of the overall continuous activity pattern and failing to reproduce the ZT12 activity peak (Fig. 2).

**Figure 2.**
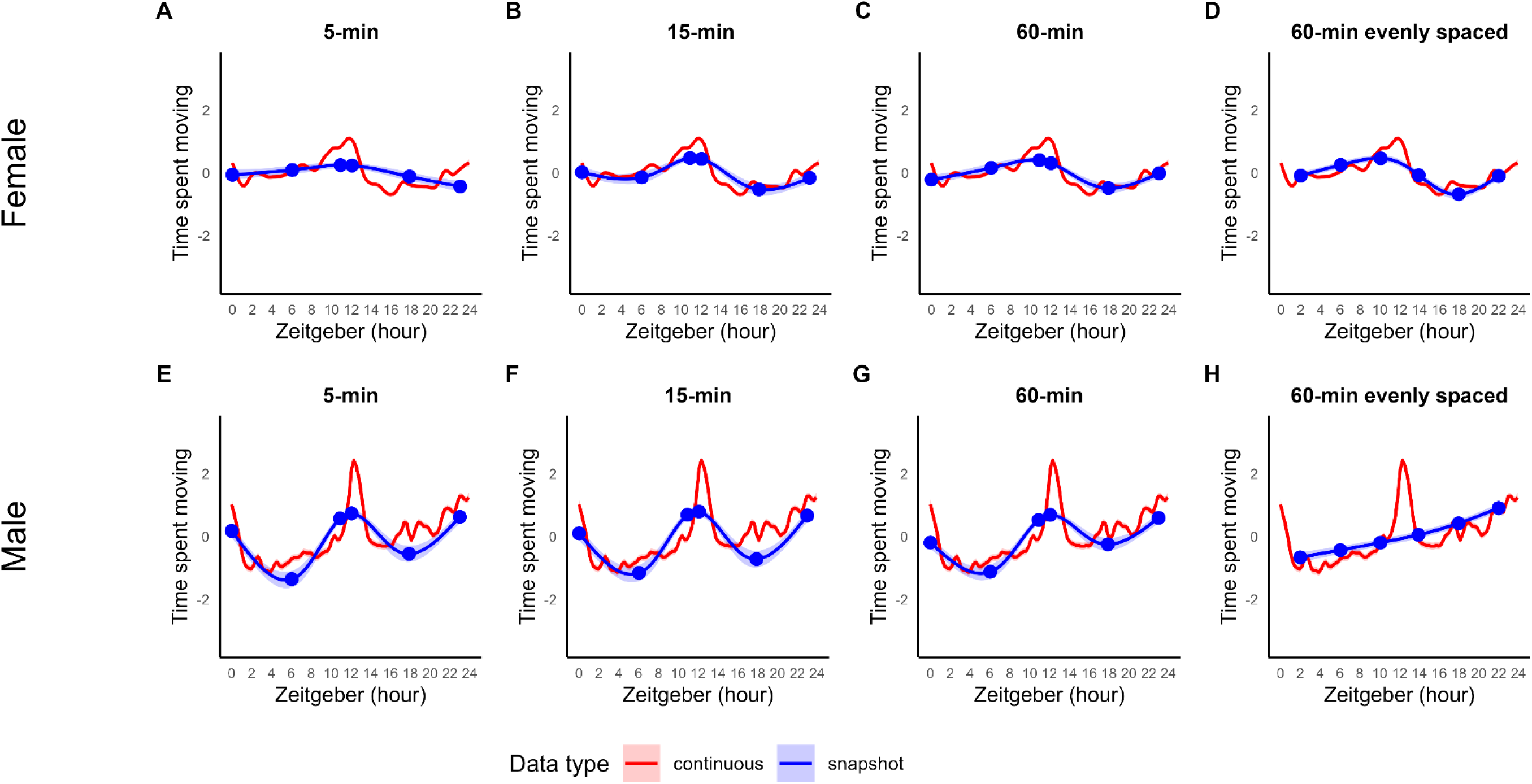
Estimated partial temporal effects for time spent moving (log scale) from continuous data (red) and snapshot data (blue), with 95% confidence intervals. Biologically informed snapshots were collected at ZT 0, 6, 11, 12, 18, 23 using observation windows of (a,e) 5-min (b,f) 15-min and (c, g) 60-min. Evenly-spaced 60-min snapshots were collected at ZT 2, 6, 10, 14, 18, and 22 (d, h). Panels (a-d) show females and panels (e-h) show males. Blue dots represent the timing of the snapshot observations.

We next assessed the effect of snapshot duration, using only the biologically informed sampling schedule and comparing observation windows of 5, 15 and 60 min. Snapshot duration strongly influenced how closely fragmented sampling reproduced the continuous activity pattern, with 60-min snapshots providing the closest representation based on RMSE and R² values (Table 4). Significant temporal variation in activity was detected for both sexes across all snapshot durations, with the exception of the 5-min snapshots for males (Table 5).

**Table 5.**
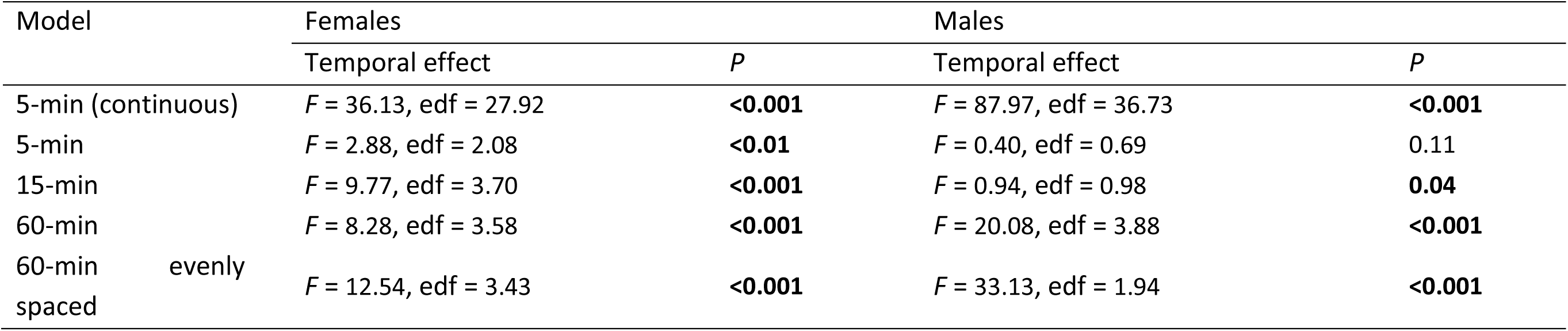
Temporal effects on time spent moving in females and males estimated using AR1 GAMM models for the continuous 5-min binned model and each of the four snapshot models (5-min, 15-min, 60-min, and 60-min of evenly spaced snapshots). Bold values represent significant p-values. The *F*-statistic and estimated degrees of freedom (edf) are reported. The AR(1) GAMM model-performance metrics are reported in Table S5.

Visual comparisons supported these quantitative patterns. The 60-min biologically informed snapshots most closely reproduced the shape of the continuous activity curves, whereas shorter observation windows progressively lost important features of the underlying temporal pattern (Fig. 2). More specifically, the timing and height of the activity peaks were not accurately captured for both males and females. Nevertheless, even the 60-min snapshots provided a less complete representation of activity than the continuous reference dataset (Figs. 1 & 2).

### 4.3 Assessing temporal aggregation and bin-size comparison

Increasing temporal aggregation (from 1 min through to 60 min bins) progressively reduced the ability of models to reproduce the fine-scale activity pattern represented by the 1-min reference data. For both sexes, models based on 5-min bins most closely reproduced the 1-min activity curves, with the lowest scaled RMSE, highest R² and strongest correlations between predicted curves (Table 6; Fig. 3). As bin size increased, scaled RMSE increased, while R² and correlations between predicted curves decline (Table 6).

**Figure 3.**
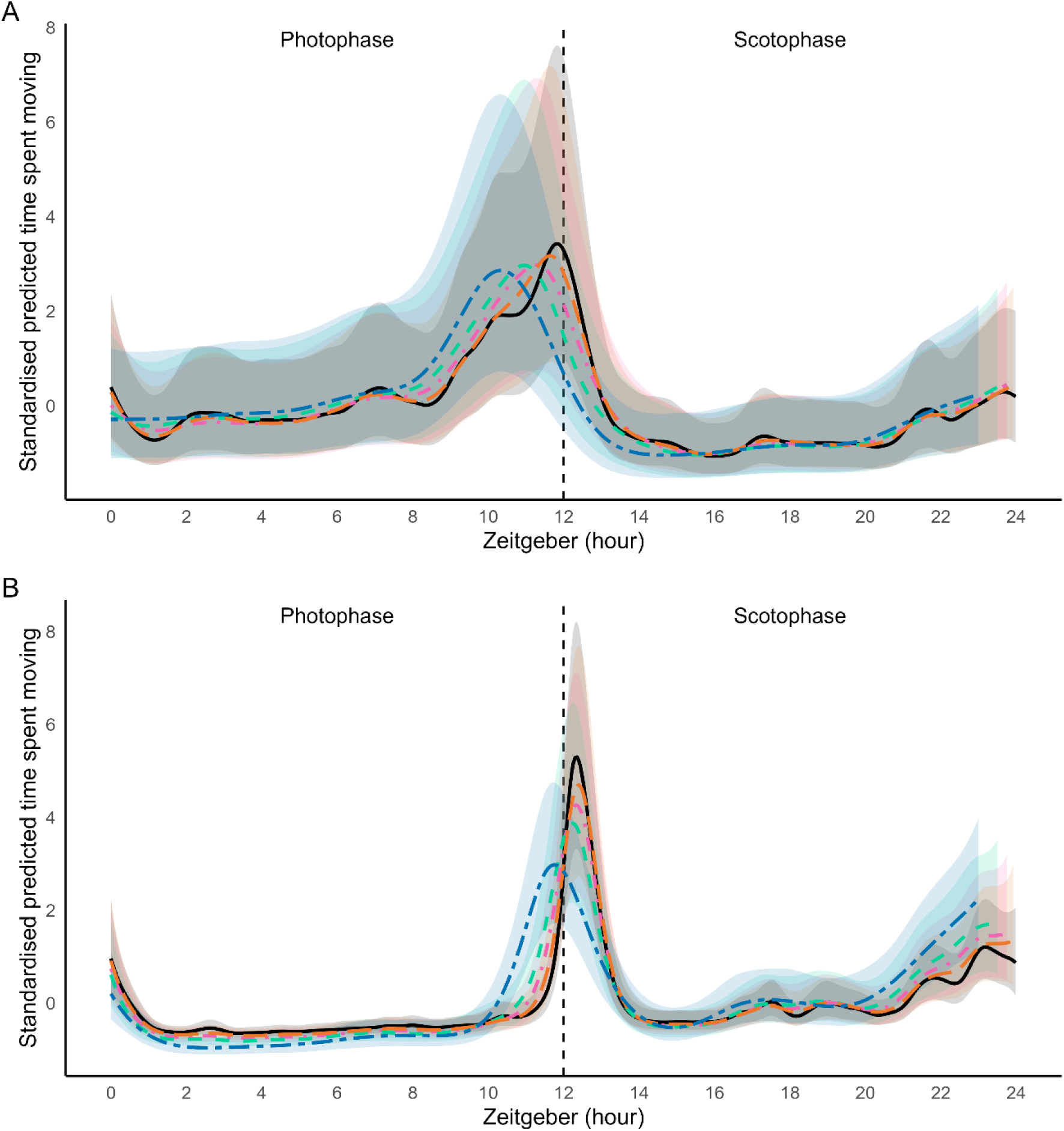
Predicted activity patterns for the time spent moving behavioural metric across temporal bin sizes over 24 h for (A) females and (B) males. Lines show predictions for 1-min (black solid); 5-min (orange dashed); 15-min (pink dash); 30-min (green dashed); and 60-min (blue dash) bins. Prediction uncertainty is represented as standardised smooth estimates with scaled confidence intervals.

**Table 6.** Similarity of activity patterns estimated from 5-, 15-, 30- and 60-min time bins to those estimated from 1-min reference data for females and males. RMSE and R² were calculated from z-standardised predictions, and correlation were calculated between predicted activity curves. Bold values indicate the closest agreement with the 1-min reference for each metric and sex.

| Bin size | Females |  |  | Males |  |  |
| --- | --- | --- | --- | --- | --- | --- |
|  | RMSE | R<sup>2</sup> | Correlation | RMSE | R<sup>2</sup> | Correlation |
| 5 min | **0.15** | **0.98** | **0.99** | **0.12** | **0.98** | **0.99** |
| 15 min | 0.24 | 0.94 | 0.97 | 0.22 | 0.95 | 0.98 |
| 30 min | 0.37 | 0.86 | 0.93 | 0.39 | 0.85 | 0.93 |
| 60 min | 0.59 | 0.65 | 0.82 | 0.77 | 0.40 | 0.73 |

Visual comparison of the predicted activity curves revealed that increasing bin size affected both the magnitude and timing of biologically important features of the activity pattern. For females, the major activity peak occurred at approximately ZT 12 in the 1-min reference model but shifted progressively with increasing aggregation, occurring at approximately ZT10 in the 60-min model (Fig. 3). The broad shape of the female activity pattern remained relatively robust, although peak magnitude decreased with increasing aggregation. For males, peak timing around ZT 12 remained comparatively robust to aggregation, but peak magnitude declined substantially (Fig. 3). At the coarsest level of aggregation (12 h), only the broad distinction between photophase and scotophase activity was retained, with both females and males showing greater activity during the scotophase (Fig. 4).

**Figure 4.**
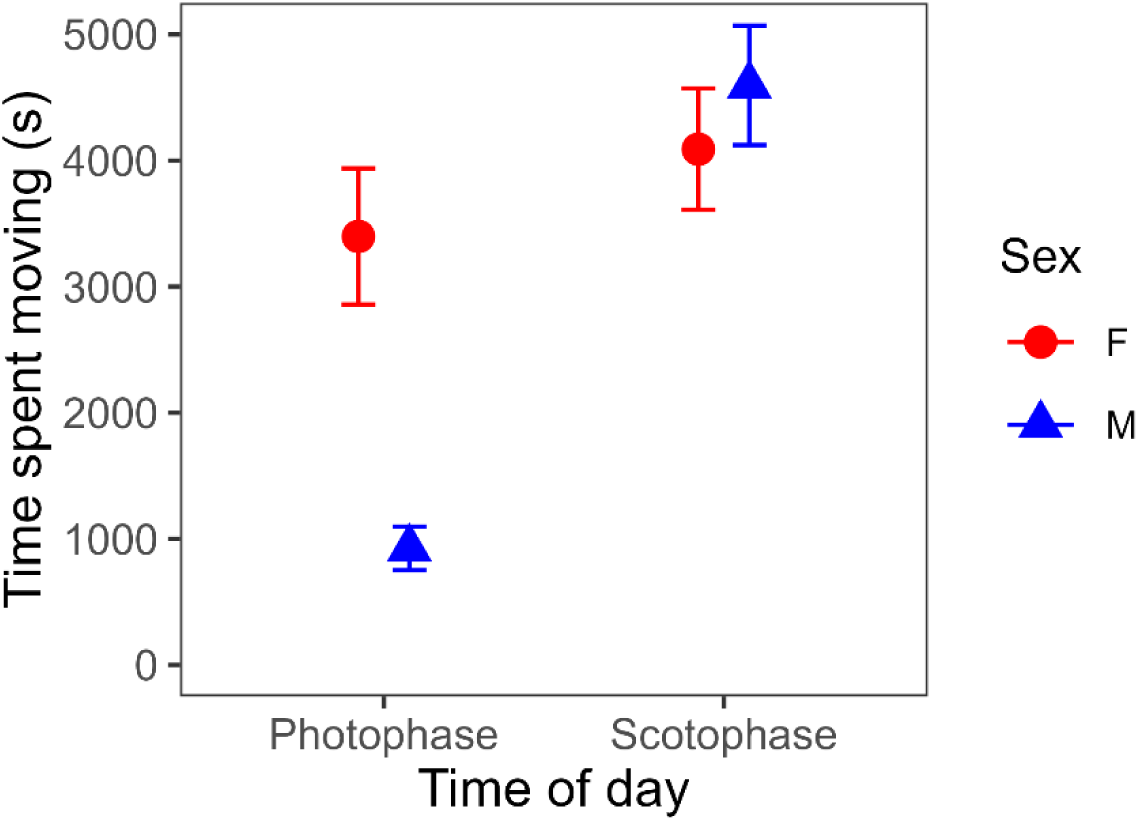
Mean (± standard error) time spent moving by females (red) and males (blue) during the 12 h photophase (ZT 0-12) and scotophase (ZT 12-24).

## 5. Discussion

Here, we demonstrate that methodological decisions made across the research workflow, from sampling design and selection of temporal resolution through to statistical model selection can shape biological inference from circadian activity data. Using high-resolution 24 h activity recordings from *D. melanogaster*, we identified three key methodological considerations relevant to the representation and analysis of circadian activity patterns. First, model choice matters: although all model classes detected temporal variation in activity, GAMMs most accurately represented the complex shape, timing and magnitude of the observed activity patterns. Second, fragmented sampling (or snapshot sampling) was not inherently inadequate, but its effectiveness depended strongly on sampling design. Longer sampling windows captured activity patterns more accurately than shorter windows, while sampling at biologically relevant times provided a better representation than evenly spaced sampling. Finally, increasing temporal aggregation progressively reduced the information retained within activity patterns, with finer bin sizes better preserving biologically important features such as the timing and magnitude of activity peaks. Importantly, our results demonstrate that statistical detection of temporal variation does not necessarily equate to accurate representation of the observed circadian activity pattern. More broadly, rather than treating model choice, sampling design and temporal resolution as independent methodological decisions, our findings show that choices across these stages collectively determine the biological information retained and ultimately inferred. We, therefore, integrate these findings into a practical decision-making framework for the collection, processing and analysis of circadian activity data (Table 8), with potential application to other behavioural time-series characterised by strong temporal structure. This framework provides a structured basis for balancing practical and analytical constraints against the need to retain sufficient temporal information for biologically meaningful inference.

**Table 7.** GAMM results for time spent moving using data aggregated into 1-, 5-, 15-, 30- and 60-min bins for females and males. Model performance is summarised by R² and percentage deviance explained (Dev. %). Temporal effects are reported as *F*-statistics with estimated degrees of freedom (edf); all temporal effects were significant (*P* < 0.0001). Age and wing size were included as covariates in all models but were not the focus of the comparison; their parameter estimates and statistical tests are reported in Table S7.

| Bin size | Females |  |  | Males |  |  |
| --- | --- | --- | --- | --- | --- | --- |
|  | R<sup>2</sup> | Dev. (%) | Female time effect | R<sup>2</sup> | Dev. (%) | Male time effect |
| 1 min | 0.25 | 23.2 | $F = 1.49$, edf = 23.89 | 0.16 | 24.1 | $F = 1.76$, edf = 31.01 |
| 5 min | 0.31 | 26.0 | $F = 1.92$, edf = 15.29 | 0.18 | 24.3 | $F = 3.38$, edf = 24.04 |
| 15 min | 0.23 | 28.7 | $F = 2.77$, edf = 14.53 | 0.18 | 28.2 | $F = 7.76$, edf = 23.00 |
| 30 min | 0.39 | 37.3 | $F = 5.37$, edf = 13.63 | 0.17 | 33.4 | $F = 13.28$, edf = 20.39 |
| 60 min | 0.38 | 39.9 | $F = 6.20$, edf = 9.91 | 0.17 | 39.1 | $F = 24.73$, edf = 14.32 |

**Table 8.** Practical decision framework for the design, processing and analysis of temporally structured behavioural data.

| Framework step | Initial approach | Evaluate | If inadequate |
| --- | --- | --- | --- |
| 1. Select model complexity | Use the simplest model appropriate to the biological question | Does the model adequately represent the temporal features required for inference? | Increase model flexibility if important peaks, asymmetry, multimodality or transitions are poorly represented |
| 2. Select sampling strategy | Use continuous sampling where feasible; otherwise use biologically informed snapshots | Do sampled periods capture the important temporal features? | Increase observation duration or temporal coverage; if timing is unknown, conduct an intensive pilot |
| 3. Select temporal resolution | Begin with fine-resolution data and progressively aggregate | Are the features required for inference retained after aggregation? | Return to a finer resolution if relevant features such as peak timing, magnitude or transitions are altered |
| Final validation: biological and environmental context | Identify the groups and conditions across which the inference will be made | Does the selected approach adequately preserve the features of interest across these contexts? | Adapt or stratify the protocol where performance differs; use pilot data where performance is unknown |

### 5.1 Model selection: how much complexity is required?

The first step in our framework is to consider what features of the temporal structure must be represented to address the biological question. Model complexity should be driven by the biological question and expected complexity of the behavioural response, rather than an assumption that either the simplest or most flexible model is inherently preferable. For relatively simple temporal responses, or where the primary aim is to detect a broad change rather than accurately characterise its shape, linear or polynomial models may provide an adequate and more parsimonious representation (Heit et al., 2024). However, their suitability decreases as temporal patterns become more complex. Linear models are poorly suited to representing nonlinear or multi-modal activity patterns, typically underperforming when compared to more complex model classes (Bera et al., 2021; de Freitas et al., 2021; Heit et al., 2024; Simpson, 2025). Additionally, for polynomial models, the global functional form can produce biologically unrealistic patterns, particularly near temporal boundaries, and increasing polynomial order can increase the risk of overfitting (Baayen & Linke, 2020; Bera et al., 2021; Simpson, 2025; Wood, 2017). Cosinor approaches serve a different purpose and may be preferable where the specific objective is to quantify rhythmicity using biologically interpretable parameters such as amplitude and acrophase (Parsons et al., 2024; Romero et al., 2010). In contrast, behaviours characterised by nonlinear, asymmetric, multi-peaked or rapidly changing temporal dynamics are likely to require greater model flexibility (Baayen & Linke, 2020; Simpson, 2025; Wood, 2025). This trade-off was evident in our *Drosophila* case study. All four model classes detected significant temporal variation, but they differed substantially in how well they represented its form. GAMMs most closely represented the shape, timing and magnitude of the observed activity patterns, whereas simpler models could detect temporal variation while misrepresenting biologically relevant features of those patterns. Thus, statistical detection of an effect should not itself be taken as evidence that a model adequately represents the underlying biology. GAMMs provide a particularly useful option where temporal structure is complex or poorly known because they can flexibly estimate non-linear response shapes without requiring a particular global function form, although this flexibility can reduce the simplicity and parameter-level interpretability offered by more constrained models (Simpson, 2025; Wood, 2017; Yee & Mitchell, 1991). Model selection should therefore balance the complexity of the biological pattern, the biological features that need to be estimated, and the costs and benefits of additional model flexibility.

### 5.2 Selecting a sampling strategy: how much and when to observe?

The second step is to determine the temporal coverage required to adequately capture the behaviour of interest. Although continuous monitoring provides the most complete representation of activity, and is considered the gold standard for capturing behaviour and activity patterns (Lynch et al., 2025; Manfrè et al., 2024; McKenzie-Smith et al., 2025; Revel et al., 2025), it may not always be feasible, particularly where logistical, technological or computational constraints limit data collection or data storage capacity (Lynch et al., 2025). Our results instead suggest that the adequacy of fragmented sampling depends on both the timing and duration of observation periods. In our *Drosophila* case study, longer snapshots more closely represented continuous activity patterns than shorter snapshots, while biologically informed sampling provided a better representation than evenly spaced sampling. Poorly timed snapshots could miss major activity peaks altogether. Consistent with this finding, Lynch et al. (2025) found that piecewise-continuous sampling can provide an effective compromise between resource-intensive continuous monitoring and simpler sampling approaches that record only whether a behaviour occurred within a sampling interval (zero-one sampling; (Brereton et al., 2022)), rather than its duration or frequency. Where continuous monitoring is not possible, the snapshot sampling should therefore prioritise observation periods of sufficient duration and at times likely to capture biologically relevant variation.

Importantly, the appropriate sampling regime is unlikely to be universal (Brereton et al., 2022; Chen et al., 2016). Where prior biological knowledge exists, sampling can be targeted towards periods when important behaviours, transitions or responses are expected (Brereton et al., 2022; Fiske & Delmolino, 2012). Where such information is unavailable, an initial pilot phase using continuous or intensive fragmented sampling could be used to characterise the temporal structure of the behaviour and identify informative sampling periods and minimum observation durations. A refined snapshot protocol could then be deployed across larger numbers of individuals, populations, sites or experimental treatments, substantially increasing sampling scale while retaining the temporal information required to address the biological question. However, even well-designed fragmented sampling cannot necessarily recover the fine-scale information contained in continuous records (Lynch et al., 2025; Manfrè et al., 2024; McKenzie-Smith et al., 2025; Revel et al., 2025). The appropriate strategy therefore depends on whether the biological question requires this fine-scale information. Sampling design should therefore be treated as an optimisation problem balancing biological coverage against the practical constraints of data collection.

### 5.3 Selecting temporal resolution: how much information?

The third step is to determine the temporal resolution required to retain the biologically relevant features of the behaviour during data processing (Tovo et al., 2016; van Leeuwen et al., 2019). As with model and sampling decisions, the smallest possible bin size should not automatically be considered optimal (Dowse & Ringo, 1994; Rajaguru et al., 2025). Instead, temporal resolution should be matched to the time scale of the behaviour and the features necessary to address the biological question (Langener et al., 2024). Behaviours sustained over relatively long periods, or studies concerned primarily with broad-scale differences in activity, may be adequately represented using larger aggregation intervals (Zhao et al., 2024), which can reduce short-term variability or noise (Dowse & Ringo, 1994), data volume and computational demands. However, where short-term variation itself contains biologically relevant information, this benefit becomes information loss: short-lived behaviours, rapid behavioural transitions, fine-scale oscillations or questions requiring accurate estimation of the timing and magnitude of activity peaks will require finer temporal resolution (Dowse & Ringo, 1994). Our *Drosophila* data illustrate this trade-off: 5-min bins closely resembled those from 1-min reference data, whereas increasing aggregation progressively reduced peak magnitude and, in females, shifted the apparent timing of peak activity. Importantly, appropriate resolution will differ among behaviours even within the same study; for example, identifying short periods of inactivity associated with sleep may require finer resolution than quantifying total distance moved over an extended period. The objective is therefore not to maximise temporal resolution indiscriminately, but to identify the coarsest level of aggregation that reduces unnecessary data complexity without losing information required for the biological inference being made (Tovo et al., 2016; van Leeuwen et al., 2019).

### 5.4 Validate the approach across relevant biological context

Importantly, decisions across all three tiers of the framework need to consider biological and individual sources of variation. The methodological approach required to adequately characterise a behaviour is not related to the nature of the behaviour itself but may depend on the individual expressing it. Relevant sources of variation may include sex (Overman et al., 2022), species (Diete et al., 2017), life-history stage (Herde & Eccard, 2013) and individual characteristics such as age (Giebultowicz & Long, 2015) or body size (White et al., 2026). Although these sources of variation were not the primary focus of our study, their inclusion revealed that methodological choices did not necessarily affect all individuals equally. Although biological variation was not the primary focus of our study, differences between females and males demonstrate that methodological choices may not affect all groups equally. For example, male activity patterns were more sensitive to fragmented sampling than female patterns: evenly spaced snapshots provided a poorer representation of the overall male activity pattern, and shorter sampling windows substantially underestimated male activity peaks. Consequently, a sampling or analytical approach that adequately represents behaviour under one biological context cannot necessarily be assumed to perform equally well under another. Biological and individual variation should therefore be considered across all three tiers when selecting model complexity, sampling design and temporal resolution. We therefore propose a final validation step following the selection of model complexity, sampling strategy and temporal resolution. Researchers should identify the biological contexts across which they intend to make inferences and access whether the approach preserves sufficiently the biological features of interest across these contexts. If performance differs, protocols may need to be adapted or stratified; where performance is unknown, pilot data across representative groups or conditions can be used to assess robustness. This validation step ensures that methodological approaches are not generalised beyond the contexts in which they have been evaluated.

### 5.5 Multidirectional application

Our framework assumes that fine-scale, high-frequency continuous data are available prior to the analysis phase and that analysis is not logistically constrained. However, many researchers face limitations in experimental logistics, computational resources, or time. In such cases, the framework can be applied in a different order, starting from data collection design rather than from an existing high-resolution dataset.

When continuous data collection is not feasible, the first step is to determine an appropriate snapshot protocol by identifying biologically informed sampling times and choosing the longest feasible observation windows. Once a snapshot scheme is established, data can be aggregated at different temporal scales and bin sizes to check for lost oscillations or noise-induced bias. Finally, the most suitable model class should be selected in light of the behaviours under study, ensuring that analytical decisions are driven by the biological questions and behavioural metrics of interest.

### 5.5 Conclusion

Our study demonstrates that biological inference from circadian and temporally structured behavioural activity depends on interconnected decisions made during data collection, processing and analysis. By evaluating model choice, sampling strategy and temporal resolution against the same high-resolution reference dataset, we show that methods can detect temporal variation while failing to preserve biologically important features of that variation. We integrate these findings into a practical decision framework in which these decisions are guided by the biological information required for inference and validated across relevant biological and environmental contexts. The overarching principle of our framework is to use the least intensive approach that demonstrably preserves the biological information needed to answer the research question. This shifts methodological choice from convention or convenience towards explicit evaluation of whether the information required for biological inference has been retained.

