## Supplementary Materials for "Capturing the rhythm: methodological choices shape inference from circadian activity data"

#### S1 Materials and Methods

##### 1.1 Bloomington's Cornmeal Recipe

Ingredients (to make 2L/ ~200 vials):

| | | |
|---|---|---|
| Boiling Water | 1100 | millilitres |
| Warm Water | 700 | millilitres |
| Tap Water | 200 | millilitres |
| Soy Flour | 40 | grams |
| Yeast | 70 | grams |
| Cornmeal | 146 | grams |
| Dextrose Monohydrate | 150 | grams |
| Agar | 12 | grams |
| Acid Mix* | 28 | millilitres |
| Tegosept | 33 | millilitres |

Method:

1. Bring large volume of water to a gentle boil in a suitably sized pot
2. Weight out and sift soy flour in a large beaker
3. Weigh yeast and cornmeal and mix them well with the soy flour
4. Weigh out dextrose then add to boiling water in a large pot
5. Weight out agar, mix with tap water (room temperature) and add to pot
6. Return to boil while stirring gently. Boil for 1 minute, stirring constantly. Turn off heat.
7. Sprinkle yeast, cornmeal and soy flour mixture into the sugar and agar solution while whisking to avoid the formation of lumps.
8. Return to the boil, stirring gently and constantly.
9. Gradually whisk in the warm water, stirring. Turn off the heat.
10. Add the acid mix and tegosept, whisking them through.
11. Dispense into bottles and vials as required, while stirring.
12. Once cool, sprinkle with yeast and plug vials.
13. Store in cold room (<3°C) for up to a month.

*Acid Mix Recipe

| | | |
|---|---|---|
| RO water | 546 | millilitres |
| Propionic Acid | 412 | millilitres |

| | | |
|---|---|---|
| Phosphoric Acid | 42 | millilitres |

Method: Add propionic acid and phosphoric acid to RO water.

#### 1.2 Filming setup

To record the exact timing of eclosion, we used eight high-resolution analogue security cameras (2 x Swann NVR8-8580 Security Camera Systems; each with 4 x NHD-875WLB cameras). These were set up on multiple covered shelves in a temperature-controlled room ($25^0$C) with a 12 h photophase (~1300 lx, cool white LED lights affixed to the shelving above) and a 12 h scotophase (< 0.01lx, accounting for minor light spill comparable to the light emitted from a new moon {Gaston, 2013 #203}). We placed between 35-40 circular experimental arenas (diameter = 29mm; vol = $4ml^3$ containing $3ml^3$ (approx. 2mm high) of grape juice agar for humidity and a thin layer of yeast paste as a food source) under each of the security cameras and added a white sheet of paper under the arenas to increase contrast. During the scotophase, we used the inbuilt infrared light sources from the video cameras to ensure individuals were visible during the dark lighting condition. Individuals were assigned randomly under each camera ensuring that all populations were represented on all shelves to reduce positioning effects. We recorded experimental arenas for up to 72 h to ensure we captured all eclosion times, with an additional 24 h to monitor activity for all flies post-eclosion.

#### 1.3 Known-age individuals

To create known-age individuals, we transferred ~300 males and females from each of the five experimental populations, to standard population-specific lay cages containing agar base plates (see Supplementary Materials) for a 24 h conditioning period {Piper, 2016 #334}. After 24hr, we replaced the agar base plates every hour for a total of six hours and transferred all agar plates (with any laid eggs) to standard lighting conditions (12:12 L:D) at 25°C and constant humidity (~65%) for 24 h to allow egg hatch. After 24 h, we collected 30 first instar larvae from each lay cage and transferred them to a fresh food vial (N = 6 replicate vials x 5 populations spanning a 6 h time-block). We left vials for four days under standard conditions (sufficient for all larvae to pupate) and then selected haphazardly five pupae from each of six replicate vials (N = 5 pupae x 6 replicate vials x 5 populations = 150 flies). Pupae were transferred to the individual circular experimental arenas and placed under the video cameras and recorded up to the timing of eclosion, which was extracted to the nearest second based on the recorded time.

#### 1.4 Activity Data Extraction

We used EthoVision XT for activity data collection (Ver. 18; Noldus Information Technology, Leesburg, {Noldus, 2002 #612}; see http://www.noldus.com/ethovision for additional references and information on EthoVision software). To reduce processing times and to ensure that activity shifts between photoperiods were identical, we extracted activity data for a 24 h period for each individual fly using a two-step approach. First, all individuals were greater than four hours old post adult eclosion. This age parameter ensured sufficient time for essential early adult behaviours; first excretion (typically 1–2 h post-eclosion), wing unfurling (within 1 h post-eclosion), and drying and melanising (3 hours post-eclosion) {Seong, 2023 #218}). Second, to standardise responses to lights on and off, and to facilitate statistical analyses we extracted activity data at the start of the first lights on phase (ZT 0) following an individual's eclosion. These two criteria meant that any adult that eclosed <4 h prior to their first lights-on (ZT 0), were placed into the following activity monitoring period resulting in adults that were aged between 4 h and 28 h post-eclosion at the commencement of their trial. To account for these differences, we included adult age, as determined by the eclosion time, in all analyses.

Prior to analysis in EthoVision XT, we converted videos to an MPEG-4 format (resolution: 1280x720 pixels and 30 fps) using a free editing software (VideoProc Vlogger, 2024). For the 12 h of scotophase in each video, we increased the contrast to 100% and brightness to 50% to improve the distinction between the *Drosophila* and the background grape juice agar. We reduced the number of sampled data points to 15 per second for faster computation within the EthoVision XT detection settings. Following data-acquisition, we used the Lowess smoothing method in EthoVision XT (averaged datapoints based on the 10 surrounding datapoints) and set a maximum distance moved between datapoints to 0.5mm (meaning any data points where the fly moved >0.5mm was recorded as a missing sample).

We then extracted the following data in 1 min bins: total distance moved (mm) (*sensu* Distance moved Center-point Total mm), the amount of time spent moving per bin (s) (*sensu* Movement Moving / Center-point Cumulative Duration s) (averaged over 3 samples with start and stop thresholds at 0.01 and 0.005 mm/s), and the frequency of movement bouts (*sensu* Movement Moving / Center-point Frequency). We extracted summary data from EthoVision XT using one-minute time bins as this was small enough to capture rest behaviour (typically considered ≥5 minutes of inactivity) but large enough to reduce noise during analysis {Chiu, 2010 #153}. To determine rest (not an analysis profile on EthoVision XT), we used the time spent moving to create an *inactivity flag* when individuals moved for less than 2% of the one-minute time bin (we determined 2% would account for any minor errors in the data collection). Using this inactivity flag, we determined that a rest bout occurred when there were five or more consecutive rest flags based on common *Drosophila* studies that define sleep (and rest as a proxy of sleep) as ≥5 minutes of inactivity {Shaw, 2000 #548} {Huber, 2004 #549;Cavanaugh, 2016 #638}. We then converted this to a binomial rest variable resulting in four metrics of circadian activity. We note that, while not an exhaustive list, the four behavioural measures capture a significant proportion of ecologically relevant circadian behaviour.

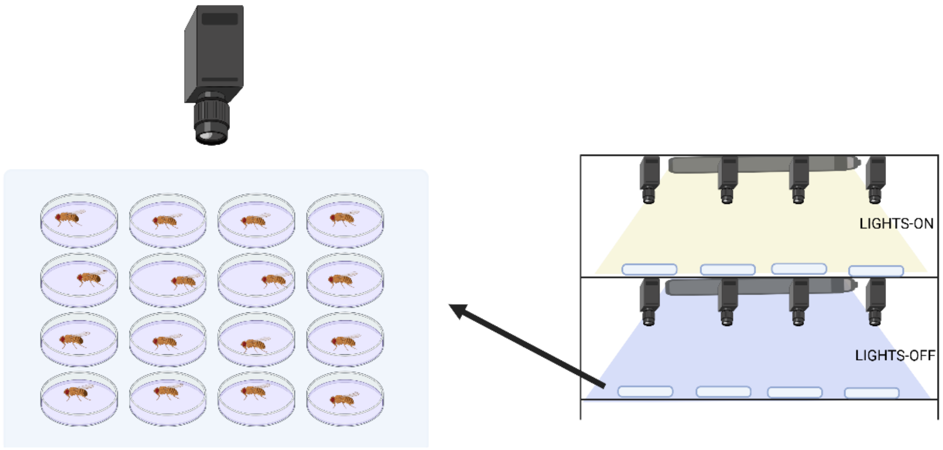


*Figure S1*. A visual representation of the filming, lighting and arena placement set up. The yellow light represents the lights-on period during ZT 0-12 and the blue light represents the lights-off period during ZT 12-24. Diagram was made using Biorender.

#### 1.4 Wing Size Length

We used wing length as an index of body size {Turner, 2011 #321}. The right wing (or left if the right was damaged) of each experimental individual was removed using a surgical blade and laid flat on a glass microscope slide. A digital image of the wing was taken under 5.12x magnification (Olympus SZX16 microscope with Olympus SDF PLAPO 1.6XPF lens). Wing length was measured from the bottom of the anterior cross vein to the point where the L3 longitudinal vein intersects the edge of the wing, using ImageJ (version 1.54d) ({Lack, 2016 #284}, Fig. S1).

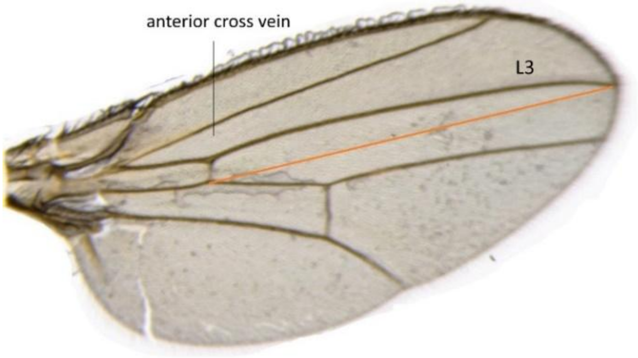

*Figure S2*. A photograph of a single *D. melanogaster* wing with the anterior cross vein and L4 vein labelled. The orange line represents the measured length from the anterior cross vein to the L4 vein which was used to determine wing size.

#### 1.5 Selected Drosophila populations and sample sizes

Table S1. Selected *Drosophila* populations that vary in their daily circadian activity or rhythmicity. The Rhythmicity Index (RI) for these populations were derived from Harbison et al (2019). Average RI values at the level of population and for each sex are presented along with the difference between males and females.

| Line | Replicate | Average RI | Male RI | Female RI |
|---|---|---|---|---|
| RI_1 | 1 | 0.09 | 0.09 | 0.09 |
| RI_2 | 2 | 0.11 | 0.09 | 0.12 |
| RI_3 | 3 | 0.12 | 0.12 | 0.11 |
| RI_4 | 4 | 0.12 | 0.13 | 0.11 |
| RI_5 | 5 | 0.12 | 0.13 | 0.11 |

Table S2. The sample sizes used in the experiment separated by replicate population and sex.

| DGRP | Replicate | Males | Females |
|---|---|---|---|
| DGRP_142 | 1 | 9 | 13 |
| DGRP_335 | 2 | 9 | 15 |
| DGRP_732 | 3 | 15 | 7 |
| DGRP_757 | 4 | 13 | 10 |
| DGRP_822 | 5 | 7 | 16 |

### S2 Data analysis

#### 2.1 General modelling overview

Table S3. An overview of the standard statistical decisions used throughout our framework analyses for all sexes and behaviours.

| Comparison | Model | Model Class | R Packages | K value | Data aggregation for model (bin size used) | Model efficacy comparison metrics |
|---|---|---|---|---|---|---|
| Model class comparison | Linear | GLMM | mgcv::bam (fREML), AR(1) via itsadug, smooth_estimates()::gratia, ggplot2 | | 1-min | Summary statistics: Akaike Information Criterion (AIC), the deviance explained, $R^2$ values, visual comparison of predicted activity curves relative to observed data |
| | Polynomial | Orthagonal and raw quartic polynomial | | | | |
| | GAMM | AR1 GAMM | | 1000 | | |
| | GLMM-cosinor | GLMM-cosinor | GLMM-cosinor | | | |
| Snapshot vs. continuous data | 5-min continuous | AR1 GAMM | mgcv::bam (fREML), AR(1) via itsadug, smooth_estimates()::gratia, ggplot2 | 200 | 5-min | RMSE and $R^2$ scores |
| | 5-min snapshot | | | 1/3 of the number of unique time points (minimum of 6 basis functions) | 5-min | |
| | 15-min snapshot | | | | 15-min | |
| | 60-min snapshot | | | | 60-min | |
| | 60-min snapshot non-biologically relevant | | | | 60-min | |
| | 1-min | AR1 GAMM | | 1000 | 1-min | |

| | | | | | | | |
|---|---|---|---|---|---|---|---|
| Bin size comparison | 5-min | | mgcv::bam (fREML), AR(1) via itsadug, smooth_estimates()::gratia, ggplot2 | based on the number of available temporal observations | 240 | 5-min | visual inspection, $R^2$ values, RMSE and correlation to reference model |
| | 15-min | | | | 94 | 15-min | |
| | 30-min | | | | 45 | 30-min | |
| | 60-min | | | | 20 | 60-min | |
| | 12 h | GLMM | glmmTMB | | | 12 h | |

### S3 Results

#### S3.1 Comparing the efficacy of various modelling approaches

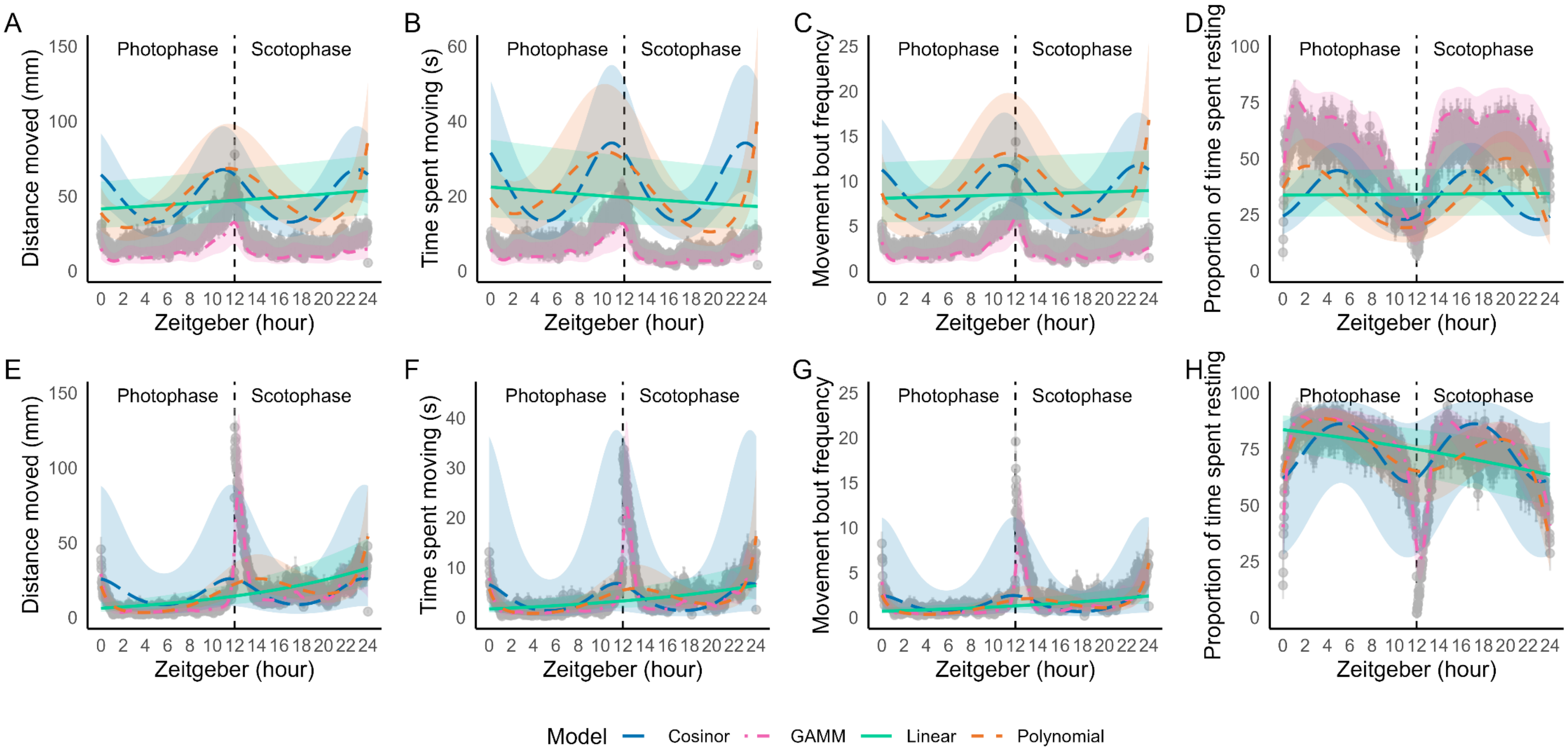


*Figure S3*. A comparison of observed and model-predicted activity patterns for (A,E) the total distance moved, (B,F) the time spent moving, (C,F) movement bout frequency and (D,H) rest behaviour, using data aggregated into 1-min bins. Panels (a-d) show females and panels (e-h) show males. Raw observed data is shown in grey (± standard errors); lines show predictions from the linear n solid), quartic polynomial (orange dashed), GLM-cosinor (blue dashed), and GAMM models (pink dashed).

Table S4A. Model-performance metrics for linear, quartic polynomial, GLMM-cosinor and generalised additive mixed models (GAMMs) fitted to the four behavioural metrics (total distance moved, time spent moving, movement bout frequency and rest) in females and males. *T*-values are presented for the age and wing size variables and *F*-statistics and estimated degrees of freedom (edf) are given for the time variable. Significant p-values and measures that indicate best fit are in bold. Bold values indicate the strongest performance and significant values ($P < 0.05$).

| | | Total distance moved | | | | Time spent moving | | | | Movement bout frequency | | | | Rest | | | |
|---|---|---|---|---|---|---|---|---|---|---|---|---|---|---|---|---|---|
| Model Type | | Linear | Polynomial (1,2,3,4) | GAMM | GLM-Cosinor | Linear | Polynomial (1,2,3,4) | GAMM | GLM-Cosinor | Linear | Polynomial (1,2,3,4) | GAMM | GLM-Cosinor | Linear | Polynomial (1,2,3,4) | GAMM | GLM-Cosinor |
| (i) Females | | | | | | | | | | | | | | | | | |
| fREML score | | 134640 | 133100 | **107810** | | 175170 | 173840 | **153000** | | 99989 | 100090 | **82600** | | 111970 | 110200 | **64484** | |
| $R^2$ | | 0.16 | 0.2 | **0.22** | | 0.19 | 0.23 | **0.25** | | 0.14 | 0.15 | **0.15** | | 0.13 | 0.17 | **0.2** | |
| Deviance explained (%) | | 20.6 | 24.2 | **26.9** | | 17 | 20.5 | **23.2** | | 11.8 | 14.6 | 16.9 | | 9.71 | 13.2 | **15.7** | |
| AIC | | 554734.1 | 550548.5 | **499055** | | 349862.1 | 346987.8 | **304178.5** | | 294169.7 | 292175.2 | **255240** | | 90427.63 | 86917.46 | **-4445.61** | |
| Time | β (SE) (edf for GAMMs) | <0.001 (<0.001) | 27.11 -1.42<br>-47.23 -1.32<br>26.26 -1.4<br>62.28 -1.3 | 31.73 | | <0.001 (<0.001) | -26.84 2.05<br>-42.87 1.86<br>57.91 1.97<br>71.35 1.89 | 23.89 | | <0.001 (<0.001) | 10.25 2.2<br>-44.88 2.03<br>28.99 2.14<br>70.24 2.01 | 27.37 | | <0.001 (<0.001) | 2.2 2.36<br>87.91 2.37<br>50.77 2.34<br>-95.22 2.36 | 20.65 | |
| | Statistic | 15.72 | 19.03<br>-35.84<br>18.81<br>47.79 | 1.31 | | -10.6 | -13.11<br>-22.99<br>29.42<br>37.8 | 1.49 | | 3.62 | 4.67<br>-22.12<br>13.54<br>34.864 | 0.86 | | 0.59 | 0.93<br>37.12<br>-21.66<br>-40.26 | 612.4 | |
| | P-value | **<0.001** | **All <0.001** | **<0.001** | | **<0.001** | **All <0.001** | **<0.001** | | **<0.001** | **All <0.001** | **<0.001** | | 0.56 | $P_1 = 0.44$<br>**$P_{2-4}$ <0.001** | **<0.000** | |
| Amplitude | β (SE) | | | | 0.3 (0.01) | | | | 0.38 (0.02) | | | | 0.3 (0.003) | | | | 0.57 (0.01) |
| | CI (5%, 95%) | | | | 0.29, 0.32 | | | | (0.35, 0.41) | | | | (0.30, 0.31) | | | | 0.55, 0.60 |
| | P-value | | | | **<0.001** | | | | **<0.001** | | | | **<0.001** | | | | **<0.001** |
| Acrophase | β (SE) | | | | -3.07 (0.03) | | | | 2.53 (0.05) | | | | 2.89 (0.01) | | | | -0.33 (0.02) |
| | CI (5%, 95%) | | | | (-3.12, -3.02) | | | | (2.43, 0.41) | | | | (2.88, 2.91) | | | | (-0.37, -0.29) |
| | Peak timing (in ZT) | | | | 11.73 | | | | 14.34 | | | | 12.96 | | | | 12.96 |
| | P-value | | | | **<0.001** | | | | **<0.001** | | | | **<0.001** | | | | **<0.001** |
| Age | B (SE) | 0.06 (0.02) | 0.07 (0.02) | 0.07 (0.02) | 0.07 (0.02) | 0.09 (0.02) | 0.09 (0.02) | 0.09 (0.02) | 0.09 (0.02) | 0.04 (0.02) | 0.04 (0.02) | 0.04 (0.02) | 0.04 (0.02) | -0.08 (0.03) | -0.08 (0.03) | -0.08 (0.03) | -0.08 (0.03) |
| | Statistic | 3.42 | 1.96 | 3.47 | (0.03, 0.10) | 3.78 | 3.78 | 3.7 | (0.04, 0.13) | 1.91 | 2.04 | 1.89 | (-0.001, 0.08) | -3.14 | -3.13 | -3.06 | (-0.13, -0.03) |

| | | | | | | | | | | | | | | | | | | | | | |
|---|---|---|---|---|---|---|---|---|---|---|---|---|---|---|---|---|---|---|---|---|---|
| | P-value | **<0.001** | **0.05** | | **<0.001** | **<0.001** | **<0.001** | **<0.001** | | **<0.001** | **<0.001** | 0.06 | **0.04** | | 0.06 | 0.06 | **<0.01** | **<0.01** | | **<0.01** | **<0.01** |
| Wing size | B (SE) | 3.78 (1.80) | 3.52 (1.80) | | 3.69 (1.80) | 3.57 (1.89) | 3.62 (2.22) | 3.42 (2.23) | | 3.57 (2.23) | 3.08 (2.35) | 3.02 (1.94) | 2.67 (1.98) | | 2.79 (2.03) | 2.74 (2.24) | -3.05 (2.16) | -3.22 (2.27) | | -3.45 (2.30) | -2.71 (2.48) |
| | Statistic | 2.1 | 3.59 | | 2.04 | (-0.14, 7.28) | 1.63 | 1.54 | | 1.61 | (-1.54, 7.69) | 1.56 | 1.35 | | 1.38 | (-1.65, 7.14) | -1.41 | -1.42 | | -1.49 | (-7.57, 2.14) |
| | P-value | **0.04** | **<0.001** | | **0.04** | **0.06** | 0.1 | 0.12 | | 0.11 | 0.19 | 0.12 | 0.18 | | 0.17 | 0.22 | 0.16 | 0.16 | | 0.13 | 0.27 |
| (ii) Males | | | | | | | | | | | | | | | | | | | | | |
| fREML score | | 136420 | 132650 | | 107690 | | 172770 | 170600 | | 147000 | | 85508 | 85654 | | 69638 | | 102410 | 101240 | | **61552** | |
| $R^2$ | | 0.1 | 0.12 | | 0.18 | | 0.09 | 0.12 | | 0.16 | | 0.12 | 0.11 | | 0.03 | | 0.11 | 0.14 | | **0.19** | |
| Deviance explained (%) | | 18.9 | 27.5 | | 36.8 | | 11.7 | 17.1 | | 24.1 | | 12.2 | 17.1 | | 22.9 | | 9.18 | 11.9 | | **15.9** | |
| AIC | | 471053.7 | 461170.2 | | 408272.8 | | 226221.8 | 223378.3 | | 178571.1 | | 194925.3 | 192514 | | 156909.5 | | 77480.28 | 75172.33 | | **-4169.89** | |
| | | | 155.3 | 1.42 | | | | 127.69 | 2.92 | | | | 116.41 | 3.08 | | | | -85.35 | 2.5 | | |
| | | | -27.1 | 1.36 | | | | -26.6 | 2.57 | | | | -13.95 | 2.86 | | | | 10.03 | 2.34 | | |
| | β (SE) | <0.001 (<0.001) | -54.58 | 1.43 | 59.74 | | <0.001 (<0.001) | -31.81 | 2.78 | 31.01 | | <0.001 (<0.001) | -32.33 | 3.04 | 33.65 | | <0.001 (<0.001) | -6.65 | 2.39 | 27.59 | |
| Time | | | 113.6 | 1.3 | | | | 136.77 | 2.6 | | | | 128.1 | 2.76 | | | | -114.33 | 2.44 | | |
| | Statistic | 85.42 | 109.5 -19.98 -38.1 87.37 | | 8.25 | | 36.17 | 43.67 -10.34 -11.43 52.66 | | 1.76 | | 30.37 | 37.78 -4.89 -10.65 46.41 | | 1.69 | | -33.72 | -34.18 4.29 -2.78 -46.85 | | 1049 | |
| | P-value | **<0.001** | **All <0.001** | | **<0.001** | | **<0.001** | **All <0.001** | | **<0.001** | | **<0.001** | **All <0.001** | | **<0.001** | | **<0.001** | **All <0.001** | | **<0.001** | |
| Amplitude | β | | | | | 0.92 (0.01) | | | | | 0.81 (0.03) | | | | | 0.44 (0.004) | | | | | 0.33 (0.01) |
| | CI (5%, 95%)) | | | | | (0.91, 0.94) | | | | | (0.76, 0.87) | | | | | (0.43, 0.44) | | | | | (0.31, 0.36) |
| | P-value | | | | | **<0.001** | | | | | **<0.001** | | | | | **<0.001** | | | | | **<0.001** |
| Acrophase | β | | | | | -1.85 (0.01) | | | | | -1.9 (0.02) | | | | | -2.01 (0.01) | | | | | 0.96 (0.04) |
| | CI (5%, 95%)) | | | | | -1.86, -1.83 | | | | | (-1.94, -1.85) | | | | | (-2.03, -2.00) | | | | | (0.89, 1.04) |
| | Peak timing (ZT) | | | | | 7.07 | | | | | 7.26 | | | | | 7.68 | | | | | 20.33 |
| | P-value | | | | | **<0.001** | | | | | **<0.001** | | | | | **<0.001** | | | | | **<0.001** |
| Age | β (SE) | 0.01 (0.01) | 0.01 (0.01) | | 0.01 (0.01) | 0.01 (0.01) | 0.01 (0.02) | 0.01 (0.02) | | 0.01 (0.02) | 0.01 (0.02) | 0.01 (0.01) | 0.01 (0.02) | | 0.01 (0.02) | 0.01 (0.01) | -0.02 (0.02) | -0.02 (0.02) | | -0.02 (0.02) | -0.02 (0.02) |
| | Statistic | 0.83 | 0.93 | | 0.59 | (-0.01, 0.04) | 0.48 | 0.56 | | 0.43 | (-0.02, 0.45) | 0.41 | 0.36 | | 0.45 | (-0.02, 0.04) | -1.46 | -1.49 | | -1.49 | (-0.05, 0.01) |
| | P-value | 0.41 | 0.351 | | 0.55 | 0.38 | 0.63 | 0.58 | | 0.67 | 0.42 | 0.68 | 0.72 | | 0.65 | 0.59 | 0.14 | 0.14 | | 0.14 | 0.11 |
| Wing size | β (SE) | 0.68 (1.59) | 0.07 (1.59) | | -0.38 (1.61) | -0.21 (1.59) | -0.66 (2.02) | -1.31 (2.12) | | -1.28 (2.20) | -1.31 (2.04) | -1.78 (1.90) | -2.4 (2.0) | | -2.39 (2.16) | -1.47 (1.9) | 1.55 (1.85) | 1.55 (1.92) | | 1.68 (2.00) | 1.3 (1.83) |
| | Statistic | 0.43 | 0.04 | | -0.23 | (-3.32, 2.90) | -0.33 | -0.36 | | -0.59 | (-5.30, 2.68) | -0.94 | -1.2 | | -1.11 | (-5.20, 2.27) | 0.82 | 0.81 | | 0.84 | (-2.30, 4.89) |
| | P-value | 0.67 | 0.97 | | 0.82 | 0.89 | 0.74 | 0.71 | | 0.56 | 0.51 | 0.35 | 0.23 | | 0.27 | 0.44 | 0.41 | 0.42 | | 0.40 | 0.48 |

S3.2 Comparing the efficacy of continuous v snapshot sampling

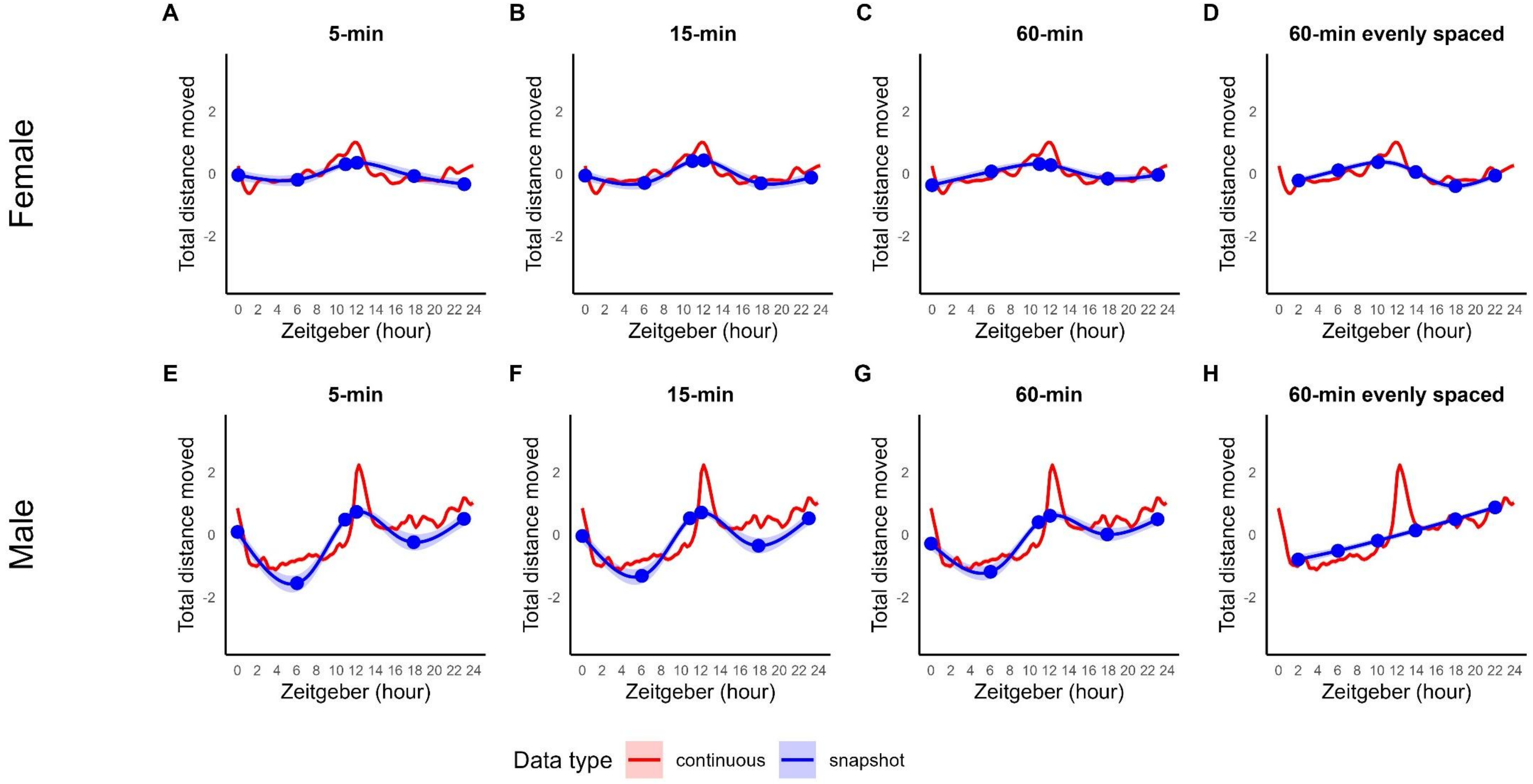


*Figure S4.* Estimated partial temporal effects for total distance moved (log scale) from continuous data (red) and snapshot data (blue), with 95% confidence intervals.

Biologically informed snapshots were collected at ZT 0, 6, 11, 12, 18, 23 using observation windows of (a,e) 5-min (b,f) 15-min and (c, g) 60-min. Evenly-spaced 60-min snapshots were collected at ZT 2, 6, 10, 14, 18, and 22 (d, h). Panels (a-d) show females and panels (e-h) show males. Blue dots represent the timing of the snapshot observations.

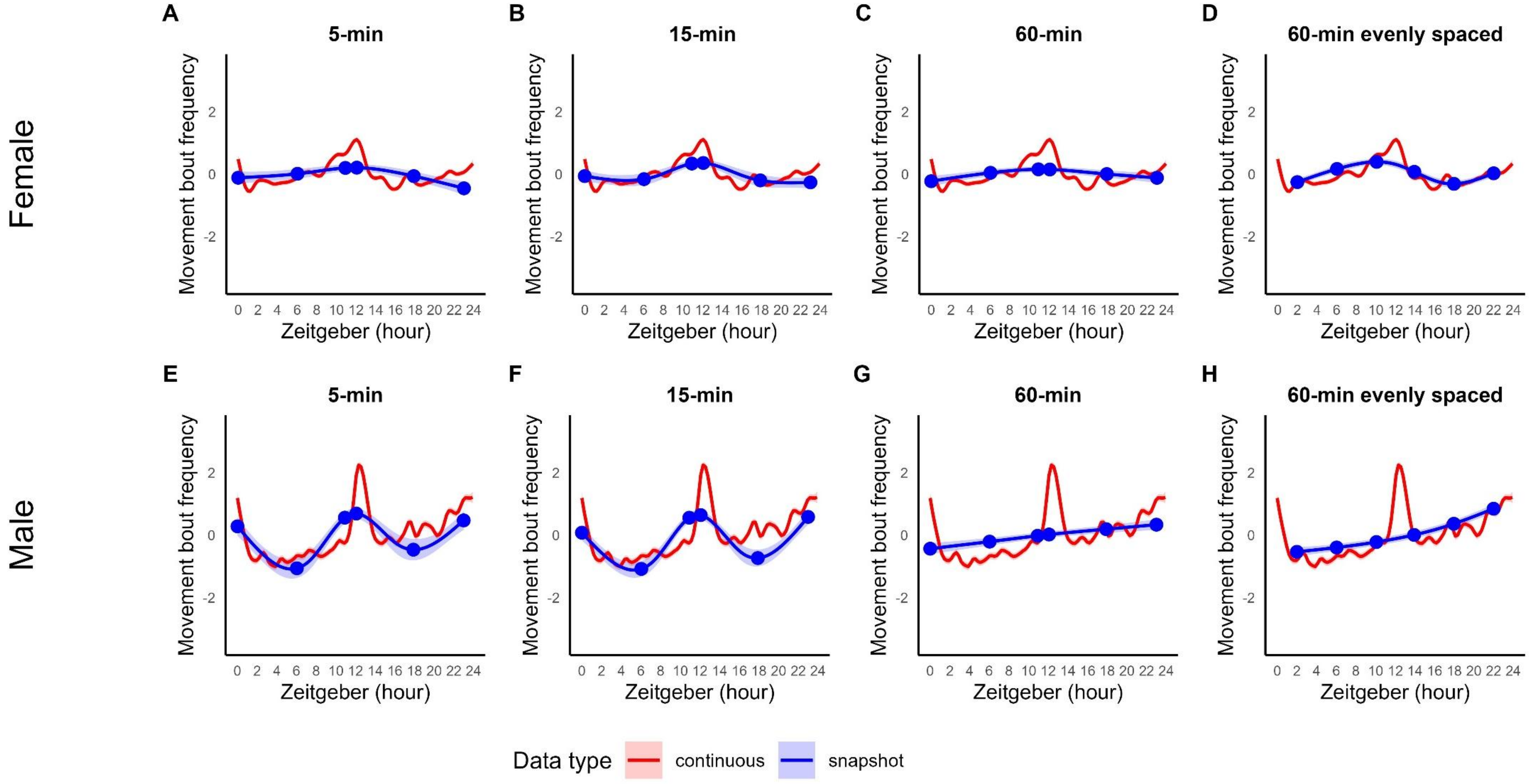

*Figure S5.* Estimated partial temporal effects for movement bout frequency (log scale) from continuous data (red) and snapshot data (blue), with 95% confidence intervals. Biologically informed snapshots were collected at ZT 0, 6, 11, 12, 18, 23 using observation windows of (a,e) 5-min (b,f) 15-min and (c, g) 60-min. Evenly-spaced 60-min snapshots were collected at ZT 2, 6, 10, 14, 18, and 22 (d, h). Panels (a-d) show females and panels (e-h) show males. Blue dots represent the timing of the snapshot observations.

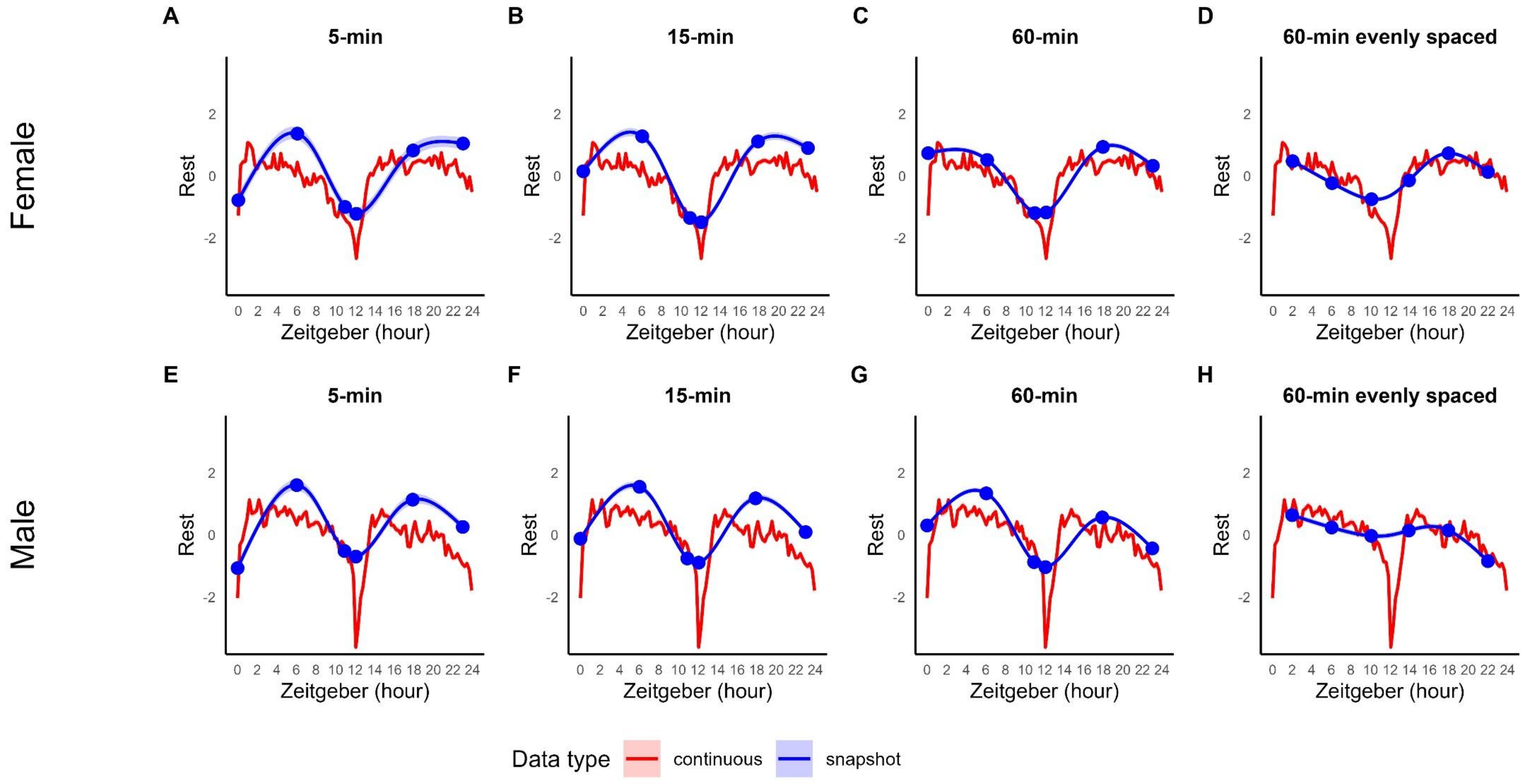


*Figure S6.* Estimated partial temporal effects for the rest behaviour (log scale) from continuous data (red) and snapshot data (blue), with 95% confidence intervals. Biologically informed snapshots were collected at ZT 0, 6, 11, 12, 18, 23 using observation windows of (a,e) 5-min (b,f) 15-min and (c, g) 60-min. Evenly-spaced 60-min snapshots were collected at ZT 2, 6, 10, 14, 18, and 22 (d, h). Panels (a-d) show females and panels (e-h) show males. Blue dots represent the timing of the snapshot observations.

Table S5. AR1 GAMM performance metrics and temporal effects for the continuous data (5-min bins) and the snapshot data (5-, 15-, 60-minutes and 60-minutes evenly spaced (ES)) for all four behavioural metrics (total distance moved, time spent moving, movement bout frequency and rest) in females and males. *T*-values are presented for the age and wing size variables and *F*-statistics and estimated degrees of freedom (edf) are given for the time variable. Significant p-values and measures that indicate best fit are in bold. Bold values indicate the strongest performance and significant values ($P$ <0.05).

| | | Total distance moved | | | | | Time spent moving | | | | | Movement frequency bout | | | | | Rest | | | | |
|---|---|---|---|---|---|---|---|---|---|---|---|---|---|---|---|---|---|---|---|---|---|
| | | Continuous data | Snapshot data | | | | Continuous data | Snapshot data | | | | Continuous data | Snapshot data | | | | Continuous data | Snapshot data | | | |
| | | 5-minutes | 5-minutes | 15-minutes | 60-minutes | 60-minutes (ES) | 5-minutes | 5-minutes | 15-minutes | 60-minutes | 60-minutes (ES) | 5-minutes | 5-minutes | 15-minutes | 60-minutes | 60-minutes (ES) | 5-minutes | 5-minutes | 15-minutes | 60-minutes | 60-minutes (ES) |
| (i) Females | | | | | | | | | | | | | | | | | | | | | |
| fREML score | | **27766** | 696 | 686 | 693 | 371 | **36440** | 770 | 772 | 795 | 467 | **22128** | 449 | 462 | 459 | 445 | **36486** | 672 | 942 | 1784 | 2024 |
| $R^2$ | | 0.27 | 0.16 | 0.19 | 0.22 | **0.35** | 0.30 | 0.18 | 0.23 | 0.27 | **0.41** | 0.17 | 0.08 | 0.11 | 0.08 | **0.27** | 0.28 | 0.32 | 0.49 | 0.53 | **0.36** |
| Deviance explained (%) | | 32.84 | 17.26 | 20.15 | 17.97 | **49.10** | 26.38 | 16.45 | 24.02 | 27.08 | **44.02** | 19.54 | 6.46 | 8.00 | 5.21 | **44.19** | 22.84 | 33.76 | 49.04 | 55.81 | **44.51** |
| Time | edf | 34.59 | 3.37 | 3.67 | 2.98 | 3.32 | 27.92 | 2.08 | 3.70 | 3.58 | 3.43 | 29.10 | 2.11 | 3.23 | 1.89 | 3.43 | 47.85 | 3.95 | 3.99 | 3.99 | 3.97 |
| | F | 89.09 | 6.09 | 8.35 | 5.06 | 10.68 | 36.13 | 2.88 | 9.77 | 8.28 | 12.54 | 27.51 | 2.42 | 4.14 | 1.64 | 10.28 | 25793.60 | 211.30 | 1026.72 | 5061.36 | 1117.78 |
| | P-value | **<0.001** | **<0.001** | **<0.001** | **<0.001** | **<0.001** | **<0.001** | **<0.001** | **<0.01** | **<0.001** | **<0.001** | **<0.001** | **0.01** | **<0.001** | **0.04** | **<0.001** | **<0.001** | **<0.001** | **<0.001** | **<0.001** | **<0.001** |
| Age | β (SE) | 0.07 (0.02) | 0.04 (0.02) | 0.044 (0.02) | 0.05 (0.02) | 0.07 (0.02) | 0.09 (0.02) | 0.05 (0.02) | 0.06 (0.02) | 0.07 (0.02) | 0.08 (0.02) | 0.04 (0.02) | 0.01 (0.02) | 0.02 (0.02) | 0.03 (0.02) | 0.05 (0.02) | -0.08 (0.03) | -0.07 (0.03) | -0.09 (0.03) | -0.08 (0.02) | -0.08 (0.03) |
| | t-value | 3.49 | 1.87 | 2.43 | 3.23 | 3.56 | 3.70 | 2.55 | 2.94 | 3.67 | 3.60 | 1.95 | 0.44 | 1.37 | 1.73 | 2.07 | -3.25 | -2.64 | -3.54 | -3.50 | -3.27 |
| | P-value | **<0.001** | 0.06 | **0.02** | **<0.001** | **<0.001** | **<0.001** | **0.01** | **<0.01** | **<0.001** | **<0.001** | 0.05 | 0.66 | 0.17 | 0.08 | **0.04** | **<0.001** | **0.01** | **<0.001** | **<0.001** | **<0.001** |
| Wing size | β (SE) | 3.84 (1.80) | 5.24 (2.29) | 5.59 (1.99) | 4.17 (1.71) | 2.66 (1.81) | 3.67 (2.26) | 4.08 (2.28) | 4.64 (2.17) | 3.16 (1.86) | 1.56 (2.24) | 2.81 (2.05) | 4.08 (2.54) | 4.47 (2.10) | 2.35 (1.66) | 1.37 (2.07) | -3.67 (2.25) | -5.66 (2.70) | -5.47 (2.41) | -2.71 (2.05) | -1.57 (2.24) |
| | t-value | 2.13 | 2.28 | 2.80 | 2.44 | 1.47 | 1.62 | 1.79 | 2.13 | 1.70 | 0.70 | 2.05 | 1.61 | 2.12 | 1.42 | 0.66 | -1.63 | -2.10 | -2.26 | -1.32 | -0.70 |

| | | | | | | | | | | | | | | | | | | | | | |
|---|---|---|---|---|---|---|---|---|---|---|---|---|---|---|---|---|---|---|---|---|---|
| | P-value | **0.03** | **0.02** | **0.01** | **0.02** | 0.14 | 0.10 | 0.07 | **0.03** | 0.09 | 0.49 | 1.37 | 0.11 | **0.03** | 0.16 | 0.51 | 0.10 | **0.04** | **0.02** | 0.19 | 0.48 |
| (ii) Males | | | | | | | | | | | | | | | | | | | | | |
| fREML score | | **25757** | 661 | 648 | 648 | 337 | **35673** | 753 | 737 | 749 | 547 | **19821** | 413 | 438 | 438 | 417 | **31666** | 743 | 1217 | 2408 | 1469 |
| $R^2$ | | **0.21** | 0.15 | 0.14 | 0.13 | 0.16 | 0.10 | 0.03 | 0.02 | **0.15** | 0.13 | -0.01 | **0.09** | **0.09** | 0.03 | 0.07 | 0.27 | 0.22 | 0.27 | **0.36** | 0.29 |
| Deviance explained (%) | | 43.18 | 16.89 | 16.68 | 15.54 | **58.45** | 24.71 | 2.65 | 2.14 | 20.81 | **37.77** | 24.48 | 9.53 | 12.33 | 4.80 | **45.27** | 24.14 | 23.58 | 32.06 | **42.29** | 41.40 |
| Time | edf | 55.82 | 3.90 | 3.89 | 3.84 | 2.29 | 36.73 | 0.69 | 0.98 | 3.88 | 1.94 | 37.37 | 3.77 | 3.86 | 1.15 | 2.00 | 88.43 | 3.96 | 3.99 | 4.00 | 3.80 |
| | f-statistic | 526.22 | 22.79 | 22.79 | 18.07 | 108.41 | 87.97 | 0.40 | 0.94 | 20.08 | 33.13 | 81.73 | 8.55 | 14.98 | 2.30 | 34.09 | 32971.19 | 183.37 | 608.21 | 3323.03 | 812.98 |
| | P-value | **<0.001** | **<0.001** | **<0.001** | **<0.001** | **<0.001** | **<0.001** | 0.11 | **0.04** | **<0.001** | **<0.001** | **<0.001** | **<0.001** | **<0.001** | **<0.001** | **<0.001** | **<0.001** | **<0.001** | **<0.001** | **<0.001** | **<0.001** |
| Age | β (SE) | 0.01 (0.01) | 0.03 (0.01) | 0.02 (0.01) | 0.01 (0.01) | 0.01 (0.01) | 0.01 (0.02) | 0.02 (0.02) | 0.02 (0.01) | 0.01 (0.01) | 0.01 (0.02) | 0.01 (0.08) | 0.02 (0.02) | 0.02 (002) | 0.01 (0.01) | 0.01 (0.02) | -0.02 (0.02) | -0.02 (0.01) | -0.03 (0.01) | -0.03 (0.01) | (-0.02 (0.02) |
| | t-value | 0.63 | 1.77 | 1.69 | 1.19 | 0.62 | 0.38 | 1.32 | 1.60 | 1.16 | 0.59 | 0.42 | 0.88 | 1.08 | 0.45 | 0.61 | -1.19 | -1.57 | -1.91 | -1.91 | -1.16 |
| | P-value | 0.53 | 0.08 | 0.09 | 0.23 | 0.53 | 0.70 | 0.19 | 0.11 | 0.25 | 0.56 | 0.68 | 0.38 | 0.28 | 0.65 | 0.54 | 0.23 | 0.12 | 0.06 | 0.06 | 0.25 |
| Wing size | β (SE) | -0.29 (1.61) | 2.15 (1.66) | 0.51 (1.62) | 0.77 (1.65) | -0.39 (1.53) | -1.58 (2.27) | 1.76 (1.88) | 1.65 (1.78) | 0.40 (1.68) | -1.98 (2.28) | -2.44 (2.18) | -1.17 (2.21) | -2.23 (1.84) | -0.35 (1.80) | -2.63 (1.17) | 1.91 (1.97) | 2.69 (1.66) | 3.25 (1.63) | 0.27 (1.76) | 2.59 (2.08) |
| | t-value | -0.18 | 1.29 | 0.31 | 0.46 | -0.25 | -0.66 | 0.94 | 0.93 | 0.24 | -0.87 | -1.12 | -0.53 | -1.21 | -0.20 | -1.21 | 0.97 | 1.62 | 2.00 | 0.16 | 1.24 |
| | P-value | 0.86 | 0.20 | 0.76 | 0.65 | 0.80 | 0.51 | 0.35 | 0.36 | 0.81 | 0.39 | 0.26 | 0.60 | 0.23 | 0.84 | 0.23 | 0.33 | 0.11 | 0.05 | 0.87 | 0.21 |

Table S6. The Root Mean Square Error (RMSE) and $R^2$ scores for the predicted continuous data from each of the four snapshot models (5-minutes, 15-minutes, 60-minutes, and 60-minutes of evenly spaced snapshots (ES)) compared to the continuous 5-minute data for all four behavioural metrics (total distance moved, time spent moving, movement bout frequency and rest). The values that indicate the best fit compared to the continuous data are in bold.

| | Total distance moved | Time spent moving | Movement bout frequency | Rest |
|---|---|---|---|---|
| (i) Females | | | | |

| Snapshot Size | 5-min | 15-min | 60-min | 60-min (ES) | 5-min | 15-min | 60-min | 60-min (ES) | 5-min | 15-min | 60-min | 60-min (ES) | 5-min | 15-min | 60-min | 60-min (ES) |
|---|---|---|---|---|---|---|---|---|---|---|---|---|---|---|---|---|
| RMSE | 0.288 | 0.239 | 0.249 | **0.232** | 0.395 | 0.274 | **0.256** | 0.273 | 0.361 | 0.285 | 0.337 | **0.266** | 0.536 | 0.621 | 0.406 | **0.363** |
| $R^2$ | 0.318 | 0.531 | 0.488 | **0.558** | 0.162 | 0.597 | **0.647** | 0.601 | 0.0805 | 0.427 | 0.199 | **0.5** | 0.162 | -0.128 | 0.518 | **0.616** |
| (ii) Males | | | | | | | | | | | | | | | | |
| Snapshot Size | 5-min | 15-min | 60-min | 60-min (NBR) | 5-min | 15-min | 60-min | 60-min (NBR) | 5-min | 15-min | 60-min | 60-min (NBR) | 5-min | 15-min | 60-min | 60-min (NBR) |
| RMSE | 0.57 | 0.576 | **0.493** | 0.561 | 0.745 | 0.715 | **0.519** | 0.638 | **0.543** | 0.598 | 0.643 | 0.614 | 0.843 | 0.926 | **0.682** | 0.697 |
| $R^2$ | 0.508 | 0.498 | **0.632** | 0.524 | 0.121 | 0.189 | **0.572** | 0.354 | **0.472** | 0.36 | 0.259 | 0.327 | -0.149 | -0.386 | **0.248** | 0.215 |

#### S3.3. Assessing temporal aggregation and bin-size comparison

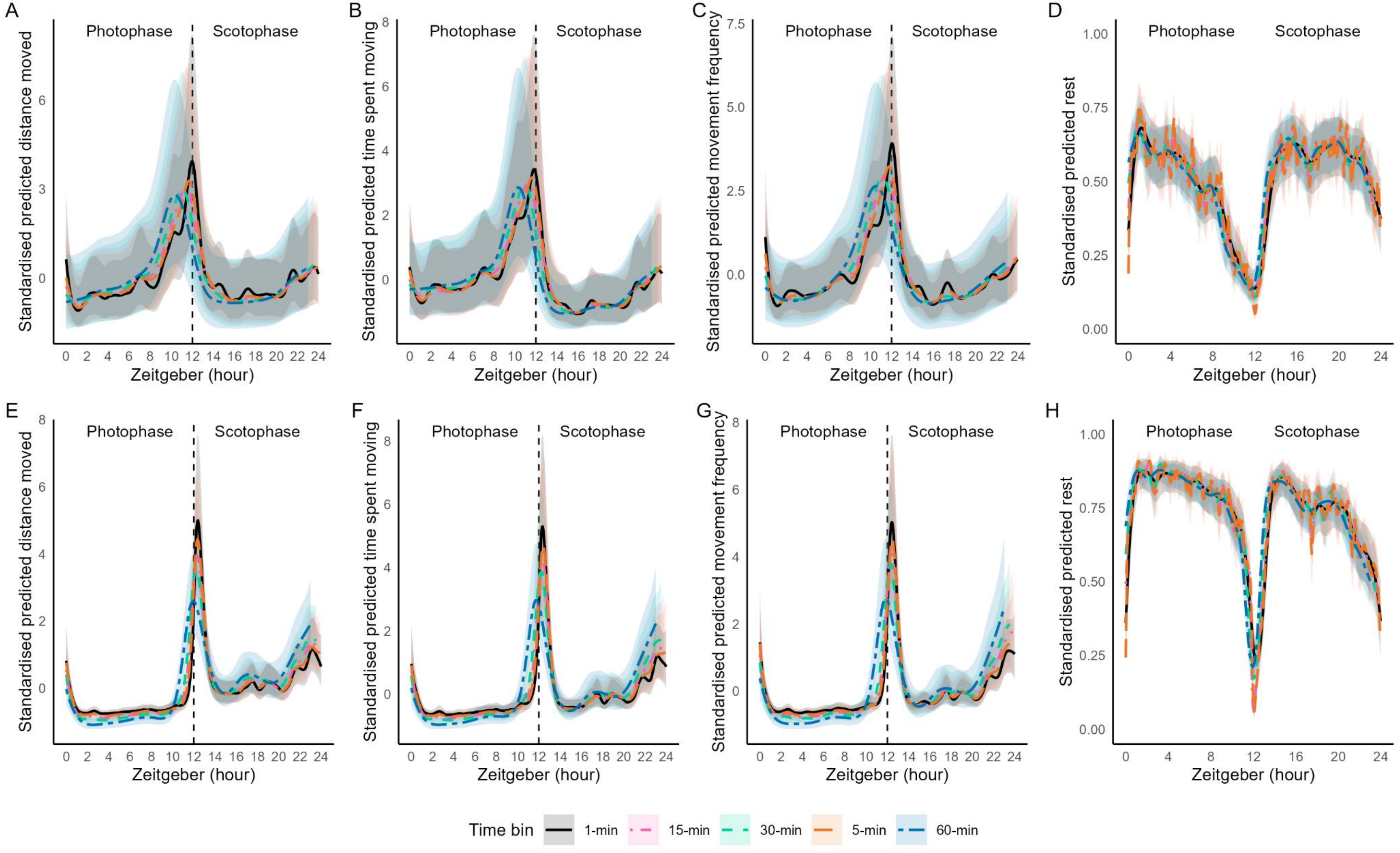


*Figure S7.* Predicted activity patterns across temporal bin sizes over 24h for (A,E) the total distance moved, (B,F) the time spent moving, (C,F) movement bout frequency and (D,H) rest behaviour. Lines show predictions for 1-min (black solid); 5-min (orange dashed); 15-min (pink dash); 30-min (green dashed); and 60-min (blue dash) bins. Prediction uncertainty is represented as standardised smooth estimates with scaled confidence intervals.

Table S7. The AR1 GAMM summary outputs for the various bin sizes (1-, 5-, 15-, 30-, 60-min and 12 h) for all four behaviours (total distance moved, time spent moving, movement bout frequency and rest). T-values are presented for the age and wing size variables and F values are given for the time variable. Significant p-values and variables that indicate best fit are in bold.

| | | Female | | | | | | Male | | | | | |
|---|---|---|---|---|---|---|---|---|---|---|---|---|---|
| | | 1-min | 5-min | 15-min | 30-min | 60-min | 12 hr | 1-min | 5-min | 15-min | 30-min | 60-min | 12 hr |
| (i) Total distance moved | | | | | | | | | | | | | |
| $R^2$ | | 0.22 | 0.28 | 0.32 | 0.34 | **0.38** | | 0.18 | 0.24 | 0.26 | 0.25 | **0.25** | |
| Deviance (%) | | 26.9 | 32.5 | 37 | 39.3 | **39.9** | | 36.8 | 42.7 | 48.2 | 50.1 | **47.7** | |
| Time | edf | 31.73 | 21.2 | 17.43 | 13.14 | 9.91 | 0.04 (0.06) | 59.74 | 36.58 | 30.76 | 24.08 | 14.82 | 1.45 (0.06) |
| | F (df) | 1.43 | 1.76 | 2.96 | 4.17 | 6.2 | 0.68 | 8.25 | 10.63 | 18.81 | 27.54 | 39.68 | 24.16 |
| | P | **<0.001** | **<0.001** | **<0.001** | **<0.001** | **<0.001** | 0.5 | **<0.001** | **<0.001** | **<0.001** | **<0.001** | **<0.001** | **<0.001** |
| Age | β (SE) | 0.07 (0.02) | 0.06 (0.02) | 0.06 (0.02) | 0.06 (0.02) | 0.06 (0.02) | 0.06 (0.02) | 0.01 (0.01) | 0.01 (0.01) | 0.01 (0.01) | 0.01 (0.01) | 0.01 (0.01) | 0.01 (0.01) |
| | T | 3.47 | 3.46 | 3.45 | 3.43 | 3.38 | 3.46 | 0.59 | 0.61 | 0.62 | 0.61 | 0.54 | 0.79 |
| | P | **<0.001** | **<0.001** | **<0.001** | **<0.001** | **<0.001** | **<0.001** | 0.55 | 0.54 | 0.53 | 0.54 | 0.59 | 0.43 |
| Wing Size | β (SE) | 3.69 (1.80) | 3.69 (1.79) | 4.05 (1.80) | 4.1 (1.80) | 4.16 (1.80) | 3.9 (1.89) | -0.38 (1.61) | -0.29 (1.60) | -0.28 (1.60) | -0.22 (01.61) | -0.22 (1.61) | -0.35 (1.62) |
| | T | 2.04 | 2.21 | 2.25 | 2.28 | 2.31 | 2.08 | -0.23 | -0.19 | -0.18 | -0.14 | -0.09 | -0.22 |
| | P | **0.04** | **0.03** | **0.02** | **0.02** | **0.02** | **0.04** | 0.82 | 0.85 | 0.86 | 0.89 | 0.93 | 0.83 |
| | | Female | | | | | | Male | | | | | |
| Bin size | | 1-min | 5-min | 15-min | 30-min | 60-min | 12 hr | 1-min | 5-min | 15-min | 30-min | 60-min | 12 hr |
| (ii) Time spent moving | | | | | | | | | | | | | |
| $R^2$ | | 0.25 | 0.31 | 0.23 | **0.39** | 0.38 | | 0.16 | **0.18** | **0.18** | 0.17 | 0.17 | |
| Deviance (%) | | 23.2 | 26 | 28.7 | 37.3 | **39.9** | | 24.1 | 24.3 | 28.2 | 33.4 | **39.1** | |
| Time | edf | 23.89 | 15.29 | 14.53 | 13.63 | 9.91 | 1.26 (0.08) | 31.01 | 24.04 | 23 | 20.39 | 14.32 | 1.26 (0.08) |
| | F (df) | 1.49 | 1.92 | 2.77 | 5.37 | 6.2 | 15.98 | 1.76 | 3.38 | 7.76 | 13.28 | 24.73 | 15.98 |
| | P | **<0.001** | **<0.001** | **<0.001** | **<0.001** | **<0.001** | **<0.001** | **<0.001** | **<0.001** | **<0.001** | **<0.001** | **<0.001** | **<0.001** |
| Age | β (SE) | 0.09 (0.02) | 0.09 (0.02) | 0.09 (0.02) | 0.09 (0.02) | 0.08 (0.02) | 0.01 (0.02) | 0.01 (0.02) | 0.01 (0.02) | 0.01 (0.02) | 0.01 (0.02) | 0.01 (0.02) | 0.01 (0.02) |
| | T | 3.7 | 3.71 | 3.7 | 3.66 | 3.63 | 0.7 | 0.43 | 0.38 | 0.36 | 0.36 | 0.31 | 0.7 |
| | P | **<0.001** | **<0.001** | **<0.001** | **<0.001** | **<0.001** | **<0.001** | 0.67 | 0.71 | 0.72 | 0.72 | 0.75 | **<0.001** |

| | | | | | | | | | | | | | |
|---|---|---|---|---|---|---|---|---|---|---|---|---|---|
| Wing Size | β (SE) | 3.57 (2.23) | 3.7 (2.21) | 3.71 (2.21) | 3.76 (2.22) | 3.71 (2.21) | -1.28 (2.05) | -1.28 (2.20) | -1.38 (2.22) | -1.49 (2.24) | -1.38 (2.24) | -1.38 (2.22) | -1.28 (2.05) |
| | T | 1.61 | 1.67 | 1.68 | 1.69 | 1.68 | -0.63 | -0.59 | -0.63 | -0.66 | -0.62 | -0.62 | -0.63 |
| | P | 0.11 | 0.09 | 0.09 | 0.09 | 0.09 | 0.53 | 0.56 | 0.53 | 0.51 | 0.54 | 0.54 | 0.53 |
| | | Female | | | | | | Male | | | | | |
| | | 1-min | 5-min | 15-min | 30-min | 60-min | 12-hr | 1-min | 5-min | 15-min | 30-min | 60-min | 12hr |
| (iii) Frequency of movement | | | | | | | | | | | | | |
| $R^2$ | | 0.15 | 0.19 | 0.22 | 0.24 | **0.26** | | 0.03 | **0.11** | **0.13** | **0.14** | 0.13 | |
| Deviance (%) | | 16.9 | 19.2 | 24.8 | 30.4 | **33.1** | | 22.9 | 24 | 29.8 | 35.5 | **29.4** | |
| Time | edf | 27.37 | 16.47 | 12.99 | 11.55 | 8.81 | 0.01 (0.06) | 33.65 | 24.46 | 23.24 | 20.67 | 14.05 | 1.24 (0.08) |
| | F (df) | 0.86 | 1.28 | 2.19 | 3.47 | 5.39 | 0.19 | 1.69 | 2.89 | 6.46 | 10.74 | 19.2 | 15.69 |
| | P | **<0.001** | **<0.001** | **<0.001** | **<0.001** | **<0.001** | 0.85 | **<0.001** | **<0.001** | **<0.001** | **<0.001** | **<0.001** | **<0.001** |
| Age | β (SE) | 0.04 (0.02) | 0.04 (0.02) | 0.04 (0.02) | 0.04 (0.02) | 0.04 (0.02) | 0.04 (0.02) | 0.01 (0.02) | 0.01 (0.02) | 0.01 (0.02) | 0.01 (0.02) | 0.01 (0.02) | 0.01 (0.02) |
| | T | 1.89 | 1.92 | 1.86 | 1.85 | 1.83 | 1.89 | 0.45 | 0.4 | 0.37 | 0.35 | 0.29 | 0.54 |
| | P | 0.06 | 0.05 | 0.06 | 1.85 | 0.07 | 0.06 | 0.65 | 0.69 | 0.71 | 0.73 | 0.77 | 0.59 |
| Wing Size | β (SE) | 2.79 (2.03) | 2.96 (2.03) | 2.51 (2.02) | 2.95 (2.02) | 3.19 (2.02) | 2.82 (2.28) | -2.39 (2.16) | -2.43 (2.15) | -2.48 (2.14) | -2.42 (2.13) | -2.48 (2.10) | -2.39 (1.92) |
| | T | 1.38 | 1.46 | 1.243 | 1.455 | 1.578 | 1.24 | -1.11 | -1.131 | -1.157 | -1.136 | -1.173 | -1.25 |
| | P | 0.17 | 0.14 | 0.21 | 0.15 | 0.12 | 0.22 | 0.27 | 0.26 | 0.25 | 0.26 | 0.24 | 0.21 |
| | | Female | | | | | | Male | | | | | |
| Bin size | | 1-min | 5-min | 15-min | 30-min | 60-min | 12-hr | 1-min | 5-min | 15-min | 30-min | 60-min | 12hr |
| (iv) % Rest | | | | | | | | | | | | | |
| $R^2$ | | 0.2 | 0.24 | 0.31 | 0.38 | **0.44** | | 0.19 | 0.24 | 0.31 | 0.37 | **0.44** | |
| Deviance (%) | | 15.7 | 20.2 | 27.6 | 34.8 | **42.1** | | 15.9 | 20.9 | 27.9 | 34.5 | **42.8** | |
| Time | edf | 20.65 | 132.48 | 78.89 | 40.41 | 18.5 | 0.24 (0.02) | 27.59 | 140.47 | 79.75 | 41.2 | 18.84 | -0.55 (0.02) |
| | χ | 612.4 | 5950 | 19948 | 15065 | 22121 | 14.46 | 1048.8 | 7990 | 15919 | 22027 | 29111 | -30.98 |
| | P | **<0.001** | **<0.001** | **<0.001** | **<0.001** | **<0.001** | **<0.001** | **<0.001** | **<0.001** | **<0.001** | **<0.001** | **<0.001** | **<0.001** |
| Age | β (SE) | -0.08 (0.03) | -0.09 (0.03) | -0.09 (0.03) | -0.09 (0.03) | -0.09 (0.03) | -0.08 (0.03) | -0.02 (0.02) | -0.03 (0.02) | -0.02 (0.017) | -0.03 (2.02) | -0.03 (0.02) | -0.02 (0.02) |
| | χ | -3.06 | -3.12 | -3.11 | -3.1 | -3.11 | -3.18 | -1.49 | -1.47 | -1.44 | -1.52 | -1.51 | -1.57 |
| | P | **<0.01** | **<0.01** | **<0.01** | **<0.01** | **<0.01** | **<0.01** | 0.14 | 0.14 | 0.15 | 0.13 | 0.13 | 0.12 |
| Wing Size | β (SE) | (-3.45 (2.30) | (-3.1 (2.25) | (-3.7 (2.36) | (-3.65 (2.35) | (-3.47 (2.34) | (-2.67 (2.40) | 1.68 (2.00) | 1.78 (0.02) | 1.81 (2.05) | 1.67 (0.02) | 1.61 (2.0) | 1.36 (1.83) |

| | | | | | | | | | | | | |
|---|---|---|---|---|---|---|---|---|---|---|---|---|
| χ | -1.49 | -1.38 | -1.56 | -1.55 | -1.48 | -1.11 | 0.84 | -0.87 | 0.88 | 0.83 | 0.8 | 0.74 |
| P | 0.13 | 0.17 | 0.12 | 0.12 | 0.14 | 0.27 | 0.4 | 0.38 | 0.38 | 0.41 | 0.42 | 0.46 |

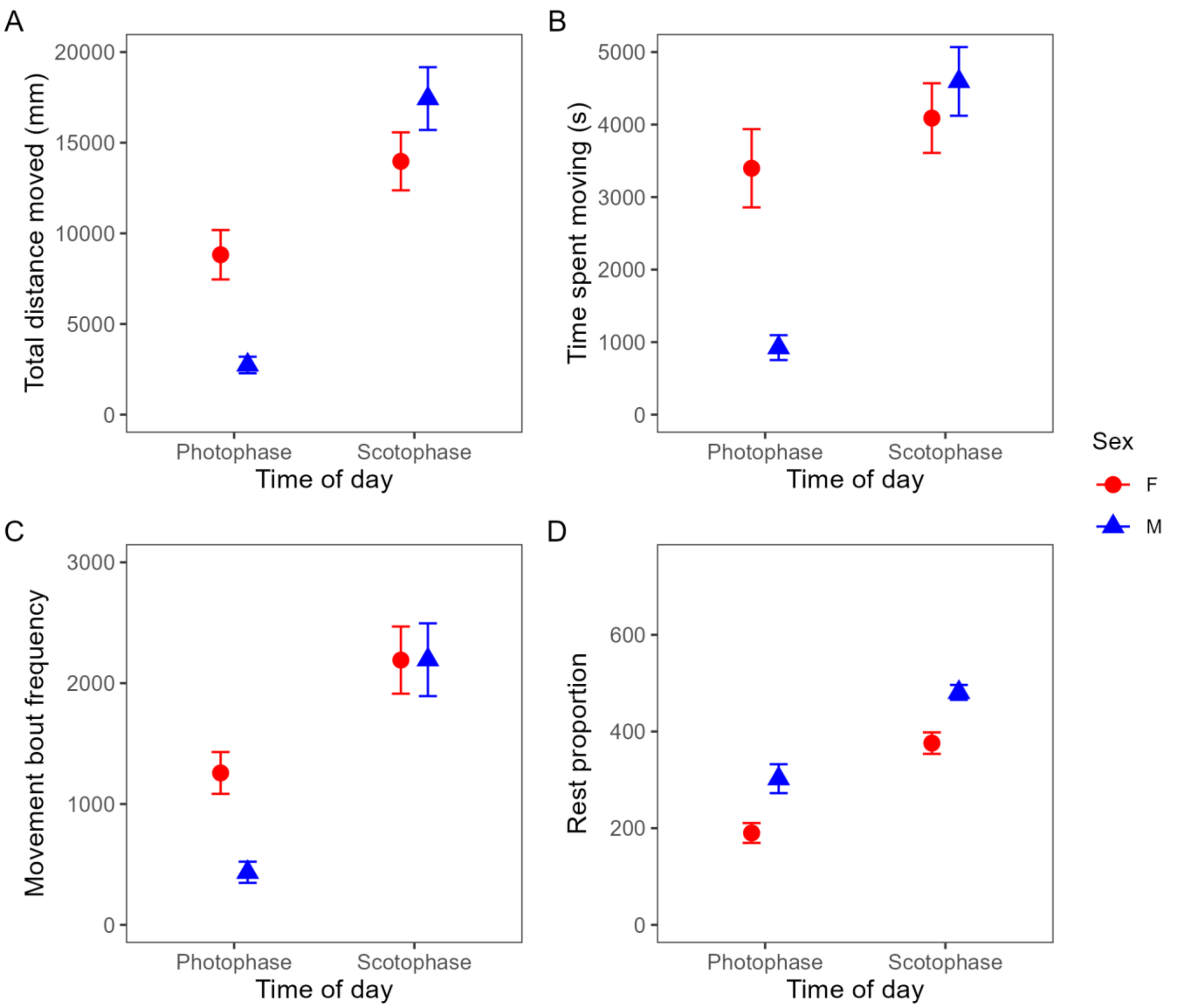


*Figure S8*. Mean (± standard error) for each of the four behaviours (A) the total distance moved, (B) the time spent moving, (C) movement bout frequency and (D) rest behaviour by females (red) and males (blue) during the 12 h photophase (ZT 0-12) and scotophase (ZT 12-24).
